# Changing reward expectation transforms spatial encoding and retrieval in the hippocampus

**DOI:** 10.1101/2020.09.13.295469

**Authors:** Seetha Krishnan, Chery Cherian, Mark. E. J. Sheffield

**Affiliations:** Department of Neurobiology and Grossman Institute for Neuroscience, Quantitative Biology and Human Behavior, University of Chicago, Chicago, IL 60637, USA

## Abstract

Internal states of reward expectation play a central role in influencing the strength of spatial memories. At the cellular level, spatial memories are represented through the firing dynamics of hippocampal place cells. However, it remains unclear how internal states of reward expectation influence place cell dynamics and exert their effects on spatial memories. Here we show that when reward expectation is altered, the same environment becomes encoded by a distinct ensemble of place cells at all locations. Further, when reward expectation is high versus low, place cells demonstrate enhanced reliability during navigation and greater stability across days at all locations within the environment. These findings reveal that when rewards are expected, neuromodulatory circuits that represent internal reward expectation support and strengthen the encoding and retrieval of spatial information by place cells at all locations that lead to reward. This enhanced spatial memory can be used to guide future decisions about which locations are most likely to lead to rewards that are crucial for survival.

## Introduction

Reward expectation is an internal state that develops through associative learning between external cues and rewards (Kobayashi and Schultz, 2014). Cues that lead to expected rewards, such as spatial locations within environments, have high survival value and are better remembered than locations that do not lead to rewards (Braun et al., 2018; Gruber et al., 2016; Healy and Jozet-Alves, 2010; Kennedy and Shapiro, 2009; McNamara et al., 2014; Singer and Frank, 2009). Reward expectation present at locations distant from actual reward sites must therefore influence the encoding and recall of those locations, but how reward expectation exerts its effects at the cellular level remains poorly understood. The hippocampus encodes and retrieves spatial memories through populations of principal neurons called place cells which fire at specific locations in an environment (Bird and Burgess, 2008; O’Keefe and Dostrovsky, 1971). The dynamics of place cells, such as their trial-to-trial reliability and stability across re-exposures (reactivation), is associated with spatial memory encoding and recall (Cacucci et al., 2007; Dupret et al., 2010; Kentros et al., 2004; Rotenberg et al., 1996; Smith and Mizumori, 2006b). Rewards themselves, and their specific locations within environments, are represented through an accumulation of place cells tuned to those locations (Danielson et al., 2016; Hollup et al., 2001; Kaufman et al., 2020; Lee et al., 2006; Mamad et al., 2017; Poucet and Hok, 2017; Tryon et al., 2017), although see (Duvelle et al., 2019; Tabuchi et al., 2003). This, however, fails to explain how reward expectation influences the encoding and recall of locations within environments that are distant from, but lead to, rewards. We therefore set out to test whether changing internal reward expectations influences spatial encoding and recall through a modulation of hippocampal place cell dynamics at locations that lead to reward.

A key issue that has prevented the influence of reward expectations on place cells from being determined is that omitting rewards for long enough to change reward expectation alters navigation behaviors which affects place cell encoding (McNaughton et al., 1983). We therefore designed an approach that measured reward expectancy in mice as they traversed a rewarded or unrewarded spatial environment, during periods of matched navigation behaviors. By imaging the dynamics of large populations of hippocampal cells during this task, we found when reward expectation is high, place cells encode all locations within environments that lead to expected rewards with high trial-by-trial reliability. These place cells are also relatively stable across days upon re-exposure to the same environment, suggesting robust memory recall. This higher stability exists at both the single cell and ensemble level. In the same animals, lowering reward expectation caused partial remapping of the place cell ensemble and diminished both reliability and stability of place cells at all locations in the environment. Notably, removing reward did not drive these changes, only when reward expectation diminished did these changes occur. We conclude that internal states of reward expectation exert their effects on spatial memories by defining which cells represent locations that lead to rewards and by modulating their reliability and stability, likely through a dopaminergic-hippocampal circuit.

## Results

Mice were trained to run on a treadmill along a 2 m virtual linear track for water rewards (rewarded-condition: R) delivered at the track end (Fig. 1A and B), after which they were teleported back to the start. Well-trained mice learned the location of the reward and pre-emptively licked before the reward location (pre-licking), providing a lap-wise behavioral signal of reward expectation (Supplementary Fig. 1) (Waelti et al., 2001). On experimental day, mice ran in R before water reward was unexpectedly removed (unrewarded-condition: UR). Interestingly, mice continued pre-licking for a few laps in UR, as though still expecting a reward. Most laps in UR matched behavior in R (Supplementary Fig. 2A; see Supplementary Fig. 3 blue laps versus orange laps). After mice traversed UR for 10 mins, reward was reintroduced (re-rewarded-condition: RR).

**Fig. 1.**
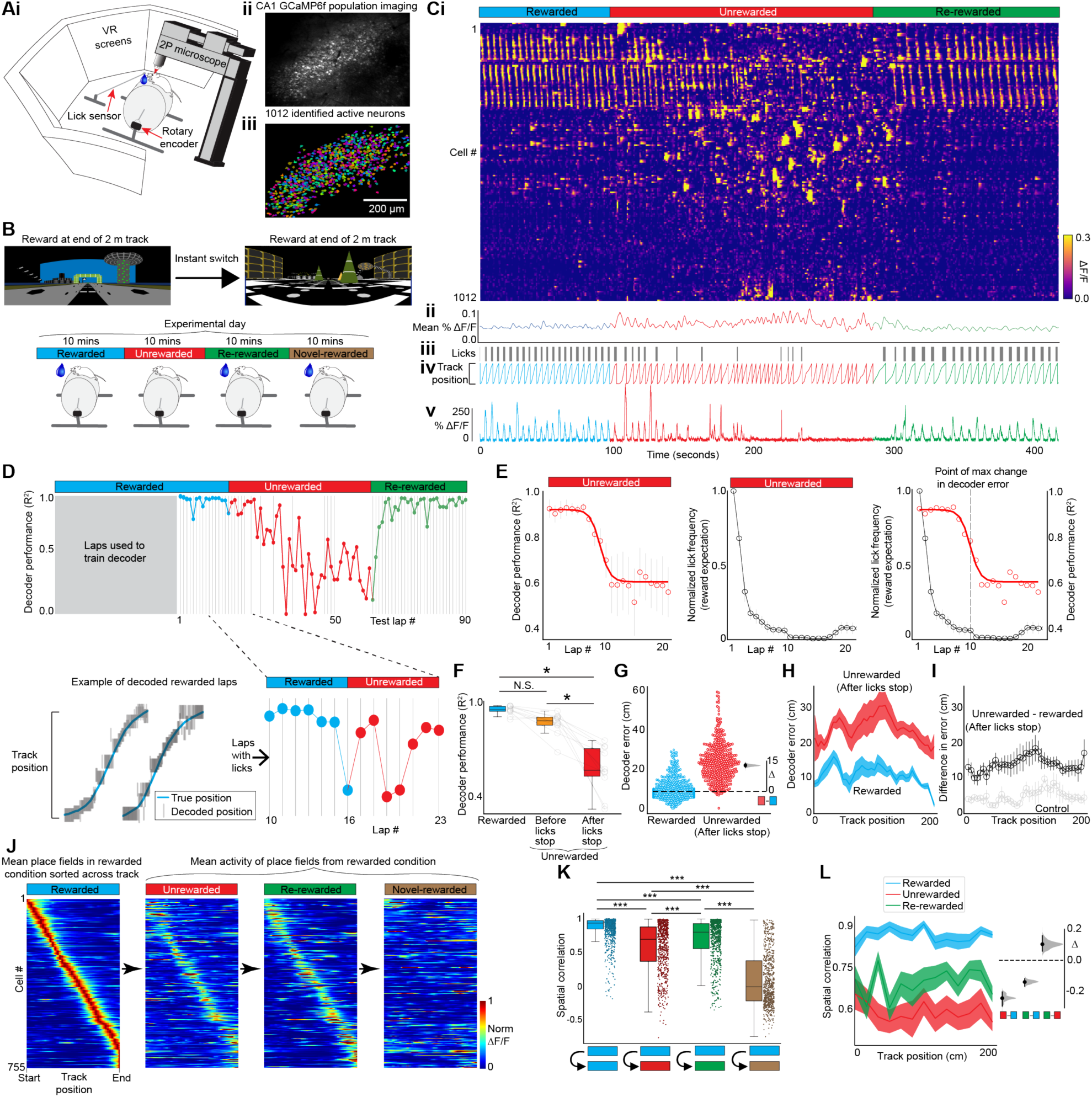
Diminished reward expectation restructures CA1 spatial encoding across the entire environment. (A) i: Experimental setup. ii: Example CA1 field of view. iii: Extracted regions of interest, randomly colored. (B) Virtual tracks (top), experimental conditions (bottom, Rewarded: R, Unrewarded: UR, Re-Rewarded: RR, Novel-rewarded: NovR). (C) i: Raster plot of ΔF/F traces from cells in Aiii arranged by correlation (highly correlated cells close together). ii: Mean ΔF/F of cells in (i). iii: Mouse licks. iv: Mouse track position. v: Example cell ΔF/F. (D) Bayesian decoder trained on initial laps activity in R. Coefficient of determination (R^2^) between true and predicted position of tested laps (top) with example fit (bottom-left). A-D are the same animal. (E) Mean lap decoder performance fitted with a reverse Boltzmann Sigmoid (r = 0.94; left) and normalized lap lick frequency (middle) in UR from all mice (n = 11). Combined (Right). (F) Mean decoder performance from all laps in R and UR. UR laps separated by when pre-licking stops. (G) Distance between true and predicted position (Decoder error: DE) at each track position. Each circle shows mean DE from a single lap pooled from all mice. Estimation plot (right) of median differences (Δ). (H) Same data, averaged by track position. (I) Difference in DE between R and UR (black), and R and R (gray; control). (J) Place fields defined in R plotted across all conditions (normalized to peak in R). (K) Mean place field spatial correlation for cells in J (dots) within R (blue) and between R and other conditions. (L) Same data, averaged by track position. Estimation plots (right). Error bars and shading indicate S.E.M. *P < 0.05 ***P < 0.001.

Using 2-photon calcium imaging of CA1 pyramidal neurons expressing the genetically encoded calcium indicator GCaMP6f (Chen et al., 2013) (Fig. 1A), we measured population activity while mice were switched across conditions: R-UR-RR (Fig. 1B and C). We found that removing reward caused a dramatic change in population activity (Fig. 1C). This was not a consequence of time (Supplementary Fig. 4) and like changes in pre-licking, did not occur immediately after reward removal (Fig. 1C). To understand this, we trained a decoder using CA1 activity during the initial laps in R and asked it to decode track position on the final laps of R and all laps in UR-RR (Methods). The decoder performed well on final laps in R and initial laps in UR before abruptly underperforming (Fig. 1D to F). Because mice pre-licked for a few laps in UR, we asked if decoder underperformance was associated with reduced reward expectation. We quantified pre-licking on each lap after reward removal and indeed found, on average, pre-licking continued for a few laps before rapidly dropping, reaching zero around the same lap decoder performance sharply dropped (Fig. 1E). This indicates that hippocampal spatial encoding remains unchanged following reward removal, until reward expectation diminishes, at which point the code abruptly transforms.

To further quantify this, we identified the lap on which pre-licking stopped in each mouse (Methods). Indeed, decoder performance in UR was similar to R until licking stopped at which point performance dropped. (Fig. 1F). This held true independent of our definition of when licking stopped (Supplementary Fig. 5 and 6). To quantify decoder error across the track, we measured the absolute distance between the true position from the predicted position at each point on the track. Interestingly, decoder error had increased across all locations within the environment that led to the reward (Fig. 1G and H), and not just around the reward site, as may be expected (Hollup et al., 2001; Kaufman et al., 2020; Lee et al., 2006; Mamad et al., 2017; Poucet and Hok, 2017). This reduced decoder performance was not explained by differences in running velocity or time (Supplementary Fig. 2 and 4; Methods). In RR, decoder performance increased, although it remained lower than in R (Supplementary Fig. 7). This provides evidence against spatial encoding being independent from reward expectations (Duvelle et al., 2019) and demonstrates that changing reward expectation drastically alters spatial encoding at all locations within an environment.

Next, we tested an alternate explanation; that inattentiveness in UR caused the changes in spatial encoding (Fenton et al., 2010; Kentros et al., 2004). We noticed that mice in R slowed down as they approached the reward site, exhibiting attention to the virtual environment (Gauthier and Tank, 2018) (Supplementary Fig. 8A). To confirm this, we exposed mice to a dark environment without any virtual cues and indeed found an absence of approach behavior (Supplementary Fig. 8A). Therefore, using approach behavior as a measure of attention, we found mice displayed attention on most laps in UR even after they stopped licking (Supplementary Fig. 8A to C). In contrast, laps displaying inattention were rare (Supplementary Fig. 8C, right). Importantly, we found the same reduction in decoder performance in UR when using only attended laps (Supplementary Fig. 8D and E). Therefore, changes in spatial encoding in UR are unlikely due to inattention but instead are due to diminished reward expectation.

The changes in spatial encoding in UR suggests place cell (PC) remapping. PC remapping is linked to encoding distinct environments (Colgin et al., 2008; Muller and Kubie, 1987; Sheffield et al., 2017). We therefore quantified remapping of PCs defined in R across UR-RR and compared to a novel rewarded environment (NovR; Fig. 1A; Methods). We focused our analysis on the period in UR when reward expectation and pre-licking had diminished (from here on UR refers only to this period). We found diminished reward expectation caused partial remapping (Fig. 1J to L), which was less than the global remapping observed in NovR (Fig. 1J and K). Partial remapping in UR occurred at all locations throughout the environment (Fig. 1L). The extent of remapping was reduced in RR (Fig. 1J to L), with some place fields (PFs) resuming their activity from R (Supplementary Fig. 9). Additionally, we found an over-representation of PFs near the reward site (Hollup et al., 2001; Kaufman et al., 2020; Lee et al., 2006; Mamad et al., 2017; Poucet and Hok, 2017) that disappeared in UR and reappeared in RR (Supplementary Fig. 10). Some of these cells we defined as “reward cells” (in addition to other cells; Supplementary Fig. 11B) (Gauthier and Tank, 2018). Reward cells showed reduced correlation in their mean activity and trial-by-trial reliability in UR (Supplementary Fig. 11C to E). Therefore, diminished reward expectation partially reorganizes the reward and place code at all locations within an environment, likely creating a distinct memory representation of the same environment (Wood et al., 2000).

Because the quality of hippocampal spatial encoding is related to memory performance (Kentros et al., 2004; Rotenberg et al., 1996), we asked whether the reorganized code in UR was of the same or reduced quality compared to the spatial code in R. We trained and tested position decoders on CA1 activity *within* each condition: R, UR, RR, and NovR (Fig. 2A to E). We found that decoder performance was lower in UR than R and RR, but similar to NovR, which could indicate that UR is encoded as a novel environment (Fig. 2A to E). We therefore measured whether spatial encoding in UR improved over time, as it does in a novel environment as it becomes familiar (Cheng and Frank, 2008; Sheffield et al., 2017). The decoder was tested on the first and last 5 laps and trained on the remaining laps (Fig. 2B). We found that decoder performance improved with time in NovR but not UR (Fig. 2D), suggesting UR is not encoded as a novel environment. Additionally, all locations in UR were poorly encoded (Fig. 2E). This shows that an environment with diminished reward expectation is encoded with an inferior spatial code at all locations that likely weakens the memory of the environment.

**Fig. 2.**
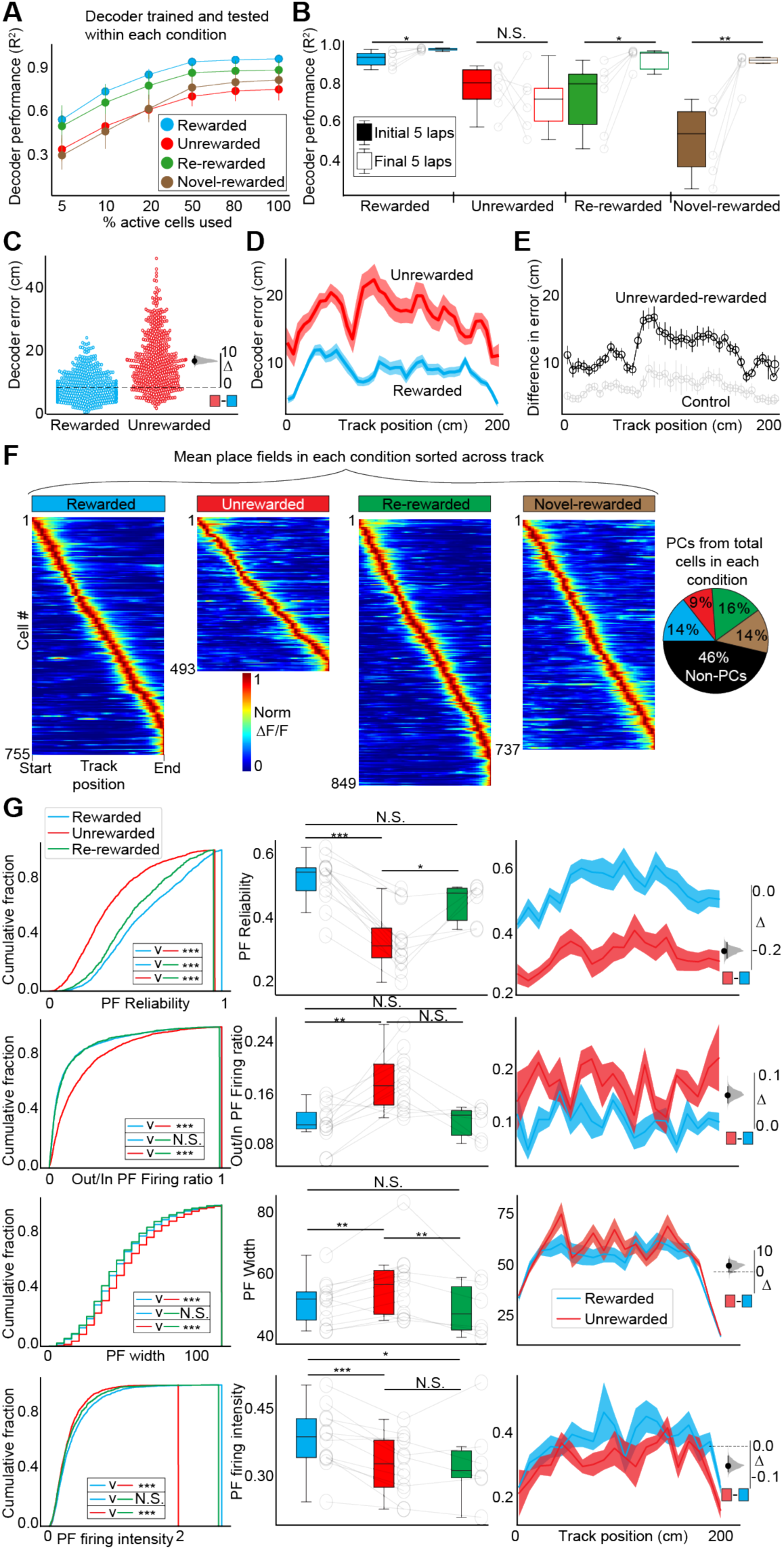
Diminished reward expectation leads to inferior spatial encoding by unreliable place cells across the entire environment. (A) Mean lap decoder performance (R^2^ of true versus predicted position) as a function of percent active cells trained and tested in each condition (6 mice). (B) Mean lap decoder performance calculated on the initial and final laps in each condition in each mouse. (C) Distance between true and predicted position (Decoder error: DE) at each track position. Each circle shows mean DE from a single lap pooled from all mice. Estimation plot (right) of median differences (Δ). (D) Same data, averaged by track position. (E) Difference in DE between R and UR (black), and between R and R (gray; control). (F) Place fields defined and sorted in each condition pooled from all mice. Each cell’s activity normalized to its peak. Pie chart shows percentage of place cells in each condition. (G) Place cell parameters are displayed as cumulative (left), average per animal (middle) and across track location (right) in R, UR and RR. Estimation plots (inset, right). Shading indicates S.E.M. *P < 0.05 **P < 0.01 ***P < 0.001.

The inferior spatial code in UR is likely due to unreliable PCs. Plotting PFs defined in each condition from all mice, we found that a reduced population of PFs encoded UR throughout the entire environment (Fig. 2F). We quantified PC properties and found that the PFs in UR had degraded on every measure of spatial encoding (Methods): PF trial-to-trial reliability (Wikenheiser and Redish, 2011), out/in PF firing ratio, PF width, and PF firing intensity, across all locations (Fig. 2G). This degradation was not time-dependent (Supplementary Fig. 12). This demonstrates that the inferior spatial code associated with diminished reward expectation is due to unreliable PCs at all locations, providing further evidence for weakened memory encoding of the environment (Kentros et al., 2004; Rotenberg et al., 1996).

Spatial memory recall is thought to be supported by the reactivation of stable PFs upon re-exposure to the same environment (Dupret et al., 2010; Kentros et al., 2004; Kinsky et al., 2018; McNamara et al., 2014; van de Ven et al., 2016). We therefore asked if PF stability across days was dependent on reward expectation by imaging the same cells across 3 days (Fig. 3A and B). Mice did not pre-lick in UR across days, demonstrating continued low reward expectation. Measuring spatial correlation of PFs across days revealed PFs in R were more correlated (Fig. 3B to G). We defined PFs as either stable or unstable across days (Methods). In each mouse, the ratio of stable/unstable PFs was significantly higher in R versus UR (Fig. 3H). Other measures of PF reliability remained reduced in UR across days similar to within sessions (Supplementary Fig. 13). Thus, at the cellular level, high reward expectation enhances PC stability across days.

**Fig. 3.**
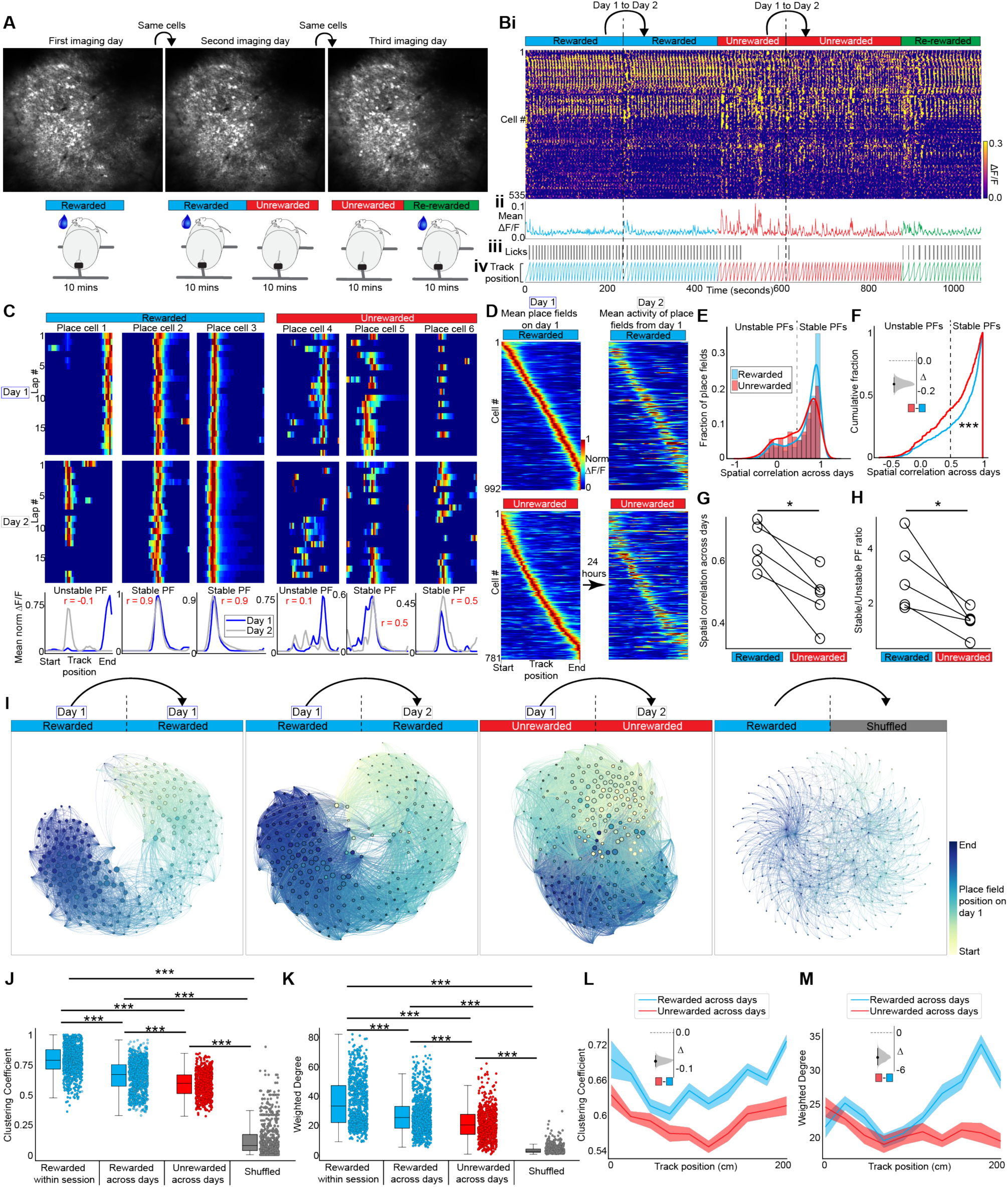
High reward expectation increases place cell stability across days at the cellular and network level. (A) Same fields of view were imaged across 3 days (n = 5 mice). (B) i: Raster plot, like Figure 1C, displaying activity of the same cells across days. ii: Mean ΔF/F from all cells. iii: Mouse behavior. (C) Example place cells across days showing lap-by-lap activity (top and middle rows; each lap normalized to its max) and their mean place fields in rewarded and unrewarded conditions (bottom). r: correlation coefficient of mean activity across days. (D) Place fields defined on Day1 plotted across days (normalized to peak on Day1). (E-G) Correlation coefficients between mean place fields on Day1 and Day2 of cells from D: (E) Histogram (F) Cumulative (G) Average per mouse. (H) Ratio of stable (correlation > 0.5) versus unstable (< 0.5) place cells across days per mouse. (I) Network graphs of neuronal activity. Nodes are place cells on Day1 colored by track position and sized by degree of connectivity. Edges correspond to correlation coefficients between Day1 and Day2. All plots are from the same mouse. Number of nodes, N, and edges, E: Rewarded Day1-Day1: N = 247, E=13132; Rewarded Day1-Day2: N=247, E=13804; Unrewarded Day1-Day2: N=245, E=15677. (J-M) Weighted degree and cluster coefficient of nodes (dots) from all animals (J and K) and averaged across track length in R and UR (L and M). Insets in F, L and M show estimation plots. *P < 0.05 ***P < 0.001.

Although the relative stability of PCs is higher when rewards are expected, many PCs are still unstable across days. However, network-level spatial encoding has been shown to be more stable across long timescales (Ziv et al., 2013). To explore if reward expectation influences network stability, we created functional network graphs from activity correlations of PCs across days in R and UR (Fig 3I, Methods). R-networks were characterized by stronger connectivity both locally (Fig. 3J) and on aggregate (Fig. 3K) than UR-networks. Local connectivity was higher among cells with nearby PFs in R than in UR across the track (Fig. 3L). Interestingly, aggregate correlations increased strikingly with proximity to reward in R but not in UR (Fig. 3M), suggesting PCs with PFs closer to reward sites are more likely to be part of a correlated ensemble across days. This is possibly driven by a ramping increase in dopamine release with proximity to reward from dopaminergic projections to the CA1(Lisman and Grace, 2005) (Supplementary Fig. 14), as such dopamine signals have been observed in other regions (Howe et al., 2013; Mohebi et al., 2019). Together, these data reveal that high levels of reward expectation stabilize spatial representations across days, which could strengthen memory recall of an environment that is expected to contain a rewarded location (Dupret et al., 2010; Gruber et al., 2016; Healy and Jozet-Alves, 2010; Howe et al., 2013; McNamara et al., 2014; Singer and Frank, 2009; van de Ven et al., 2016; Wood et al., 2000).

## Discussion

During wakeful exploration animals continuously experience external events, some of which are robustly encoded into memory for future recall. A key aspect of whether external events become encoded depends on the internal state of the animal during encoding (Tarder-Stoll et al., 2020). This work reveals that the robustness of spatial memory encoding and recall in the hippocampus is based on prior experience that sets the animal’s internal state of reward expectation. We found, at the cellular level, reward expectation modulates the structure, reliability and stability of spatial representations encoded by place cells at all locations that lead to reward (Supplementary Fig. 14). Our observations support a predictive code in the hippocampus that is strongly driven by internal reward expectations (Stachenfeld et al., 2017). Specifically, when reward expectation is high, each location in the environment leading to the reward is greatly valued because it predicts future reward (Foster and Wilson, 2006). High value locations increase the reliability and stability of place cells creating a robust memory of those locations. These results provide strong evidence that spatial representations of environments are not value-free as has been suggested (Duvelle et al., 2019; Tabuchi et al., 2003), but instead value determines the structure and dynamics of spatial representations in CA1.

Reward contingencies in navigation tasks have been shown to modulate place cells (Danielson et al., 2016; Gauthier and Tank, 2018; Lee et al., 2012; Lee et al., 2006; Lee et al., 2019; Martig and Mizumori, 2011; Tryon et al., 2017; Wikenheiser and Redish, 2011). Most studies in this area either alter reward magnitudes or move reward locations, and many times include a decision-making component in their behavioral task. These factors all modulate place cells in some way, depending on the specifics of the experiment. What has been difficult to achieve in this research area is a complete removal of rewards for long enough to alter reward expectation yet maintain matched navigation behavior. This is a necessary step to assess the influence of changing reward expectations on place cells without confounds caused by changes in behavior (McNaughton et al., 1983).

Our behavioral set-up allowed us to match navigation behaviors, even when reward was not expected. Specifically, head direction, location occupancy, location sequences leading to reward, and running speed were the same in rewarded and unrewarded conditions. A number of factors led to this matched behavior: 1) Mice were head-fixed; 2) The behavior was simple and stereotyped (mice run on a linear treadmill along a linear track); 3) Mice were first trained to run to a very high level with reward before reward was removed; 4) Many traversals of the environment could be achieved in short succession (∼5 traversals/min). In conjunction with our ability to measure reward expectation on a trial-by-trial basis, this matched behavior allowed us to specifically connect the influence of reward expectation on place cells in real-time.

Interestingly, our data show that the presence/absence of reward itself seems to have little influence on spatial encoding in the hippocampus. Only after animals learn to associate or disassociate reward from the environment and change their reward expectation levels accordingly do we observe changes in spatial encoding. The structure of the map, place cell over-representation of the reward location, and trial-by-trial reliability were all modulated only when reward expectation changed, not when reward was removed/added. This demonstrates that the act of reward attainment does not in itself modulate place cells in the hippocampus, which may have been the case through a reward-related feedback signal. Instead, the animal’s internal state of reward expectation is a stronger driver of place cell encoding than the external reward. Reward cells in this region have recently been described that encode reward independently of space (Gauthier and Tank, 2018). We did identify these cells, but found that they too were modulated by reward expectation rather than reward *per se*. Therefore, our data suggests internal states of reward expectation rather than reward attainment modulate hippocampal spatial encoding of locations within environments.

Attention is also known to modulate place cells (Fenton et al., 2010; Kentros et al., 2004). Because changes in reward expectation can alter attention, it is difficult to disentangle which factor is modulating place cells (Bourgeois et al., 2016; Maunsell, 2004). Here we were able to show that mice continued to pay attention to their position in the VR environment even when reward expectation dropped, suggesting reward expectation changes were solely behind the place cell changes we observed. However, this does not mean that attention levels were the same in the rewarded and unrewarded conditions. It is possible that although mice remain aware of their position, they were not actively attending to the environment in the same way as when rewards were expected (van Gaal and Fahrenfort, 2008). Nevertheless, we can rule out that the changes we observed when reward expectation diminished were due to inattention.

It is revealing that the CA1 can create 2 distinct representations of the same environment, one when rewards are expected, and one when rewards are not expected. This process occurred within a single session and was not observed when reward expectation was held constant throughout the session. Because place cell ensembles are thought to represent memories of environments, this result suggests the CA1 can create a distinct memory of the same environment. Changing task demands within an environment has been shown to have a similar affect (Shapiro et al., 2006; Wood et al., 2000). In addition, even when reward was put back and expectation returned to high levels within the session, spatial representations did not return to their original state. The process of lowering reward expectation and then returning to original levels led to a greater distinction between representations than when reward expectation remained constant throughout the session. This suggests the CA1 chunks external events into distinct episodes based on changes in internal expectations, even when those external events remain the same. In other words, when internal expectations are constant, unchanging external events are encoded as a single episode. When internal expectations change and then return back to original levels, unchanging external events are encoded as distinct episodes. This encoding of episodic information within the CA1 network is consistent with its proposed role in episodic memory (Smith and Mizumori, 2006a).

We observed higher place cell stability across days upon re-exposure to the environment at the cellular and network level when reward expectation was high. This occurred at all locations within the environment that led to reward, and not just around the reward site. This suggests that spatial representations are better retrieved when rewards are expected, supporting memory recall of locations that lead to expected rewards (Dupret et al., 2010; Gruber et al., 2016; Healy and Jozet-Alves, 2010; Howe et al., 2013; McNamara et al., 2014; Singer and Frank, 2009; van de Ven et al., 2016; Wood et al., 2000). Interestingly, we observed a ramping increase in the correlation of ensembles of place fields across days with proximity to reward that looks remarkably similar to ramping dopamine signals seen in other brain regions (Fig. 3M and Supplementary Fig. 14) (Howe et al., 2013; Mohebi et al., 2019). Although a dopamine ramp means the level of dopamine is not equal across the track, attractor-like dynamics could ensure dopamine influences place cells at all locations. On the basis of the known connectivity of CA1 neurons, this may arise from local inhibition within CA1 or driven by input from CA3, which does have recurrent connectivity to support attractor dynamics and also receives dopaminergic input (Colgin et al., 2010; Gasbarri et al., 1994; Gonzalez et al., 2019; Knierim and Zhang, 2012; Lisman and Grace, 2005; Rolls, 2007).

It remains unknown which specific mechanisms drive the place cell changes described here. Potentially, dopaminergic projections from ventral tegmental area (VTA) that carry information regarding the level of reward expectation within an environment could modulate synaptic transmission and plasticity in the hippocampus (Atherton et al., 2015; Engelhard et al., 2019; Frey et al., 1990; Ghanbarian and Motamedi, 2013; Howe et al., 2013; Huang and Kandel, 1995; Martig and Mizumori, 2011; Mohebi et al., 2019; Rosen et al., 2015; Tritsch and Sabatini, 2012) (Supplementary Fig. 14). Specifically, dopamine could promote glutamatergic synaptic transmission of specific sets of inputs that drive place cell firing in real-time, thus promoting lap-by-lap place field reliability (Martig and Mizumori, 2011; Tritsch and Sabatini, 2012). Dopamine could also facilitate long-term strengthening of the same synapses (Ghanbarian and Motamedi, 2013; Huang and Kandel, 1995; Rosen et al., 2015) to support the stability of place fields across days (Kentros et al., 2004). When reward expectation diminishes, dopamine release would be reduced (Howe et al., 2013; Mohebi et al., 2019). This would lead to a rapid disruption of glutamatergic synaptic transmission (Tritsch and Sabatini, 2012), triggering partial remapping and causing place cells in the new map to become unreliable at all locations in the environment (Gill and Mizumori, 2006; Mamad et al., 2017; Martig and Mizumori, 2011). The loss of dopamine would also result in a loss of facilitation of synaptic potentiation, leading to reduced place field stability across days (Martig and Mizumori, 2011; McNamara et al., 2014; Retailleau and Morris, 2018; Rosen et al., 2015). Indeed, supporting this framework, a loss of dopamine function has been shown to diminish reward-place associations (McNamara et al., 2014). Future studies can untangle the precise mechanisms that modulate place cells during changes in reward expectations that we have uncovered here. Lastly, we emphasize that these observations are a striking example of how internal states shape the way the external world is encoded and recalled by the brain to determine subjective experience.

## Materials and Methods

### Subjects

All experimental and surgical procedures were in accordance with the University of Chicago Animal Care and Use Committee guidelines. For this study, we used 10-12 week old male C57BL/6J wildtype (WT) mice (23-33g). Male mice were used over female mice due to the size and weight of the headplates (9.1 mm x 31.7 mm, ∼2g) which were difficult to firmly attach on smaller female skulls. Mice were individually housed in a reverse 12 hr light/dark cycle and behavioral experiments were conducted during the animal’s dark cycle.

### Mouse surgery and viral injections

Mice were anaesthetized (∼1%-2% isoflurane) and injected with 0.5 ml of saline (intraperitoneal injection) and 0.5 ml of Meloxicam (1-2 mg/kg, subcutaneous injection) before being weighed and mounted onto a stereotaxic surgical station (David Kopf Instruments). A small craniotomy (1-1.5 mm diameter) was made over the hippocampus (1.7 mm lateral, −2.3 mm caudal of Bregma). For population imaging, a genetically-encoded calcium indicator, AAV1-CamKII-GCaMP6f (pENN.AAV.CamKII.GCaMP6f.WPRE.SV40 was a gift from James M. Wilson – Addgene viral prep #100834-AAV1; http;//n2t.net/addgene:100834; RRID:Addgene_100834) was injected (∼50 nL at a depth of 1.25 mm below the surface of the dura) using a beveled glass micropipette leading to GCaMP6f expression in a large population of CA1 pyramidal cells. Afterwards, the site was covered up using dental cement (Metabond, Parkell Corporation) and a metal head-plate (9.1 mm x 31.7 mm, Atlas Tool and Die Works) was also attached to the skull with the cement. Mice were separated into individual cages and water restriction began the following day (0.8-1.0 ml per day). Around 7 days later, mice underwent another surgery to implant a hippocampal window as previously described(Dombeck et al., 2010). Following implantation, the head-plate was reattached with the addition of a head-ring cemented on top of the head-plate which was used to house the microscope objective and block out ambient light. Post-surgery mice were given 1-2 ml of water/day for 3 days to enhance recovery before returning to the reduced water schedule (0.8-1.0 ml/day). Expression of GCaMP6f reached a somewhat steady state ∼20 days after the virus was injected.

### Behavior and Virtual Reality

Our virtual reality (VR) and treadmill setup was designed similar to previously described setups(Heys et al., 2014; Sheffield et al., 2017). The virtual environments that the mice navigated through were created using VIRMEn(Aronov et al., 2017). 2 m linear tracks rich in visual cues were created that evoked numerous place fields in mice as they moved along the track at all locations (Fig. 1B)(Bourboulou et al., 2019). Mice were head restrained with their limbs comfortably resting on a freely rotating styrofoam wheel (‘treadmill’). Movement of the wheel caused movement in VR by using a rotatory encoder to detect treadmill rotations and feed this information into our VR computer, as in(Heys et al., 2014; Sheffield et al., 2017). Mice received a water reward (4 µl) through a waterspout upon completing each traversal of the track (a lap), which was associated with a clicking sound from the solenoid. Licking was monitored by a capacitive sensor attached to the waterspout. Upon receiving the water reward, a short VR pause of 1.5 s was implemented to allow for water consumption and to help distinguish laps from one another rather than them being continuous. Mice were then virtually teleported back to the beginning of the track and could begin a new traversal. Mouse behaviors (running velocity, track position, reward delivery, and licking) were collected using a PicoScope Oscilloscope (PICO4824, Pico Technology). Behavioral training to navigate the virtual environment began 4-7 days after window implantation (∼30 minutes per day) and continued until mice reached > 4 laps per minute, which took 10-14 days (although some mice never reached this level). This high level of training was necessary to ensure mice continued to traverse the track similarly after reward was removed from the environment. Initial experiments showed that mice that failed to reach this criterion typically did not traverse the track as consistently without reward. Such mice were not used for imaging. In mice that reached criteria, imaging commenced the following day.

### Two-photon imaging

Imaging was done using a laser scanning two-photon microscope (Neurolabware). Using a 8 kHz resonant scanner, images were collected at a frame rate of 30 Hz with bidirectional scanning through a 16x/0.8 NA/3 mm WD water immersion objective (MRP07220, Nikon). GCaMP6f was excited at 920 nm with a femtosecond-pulsed two photon laser (Insight DS+Dual, Spectra-Physics) and emitted fluorescence was collected using a GaAsP PMT (H11706, Hamamatsu). The average power of the laser measured after the objective ranged between 50-70 mW. A single imaging field of view (FOV) between 400-700 µm equally in the *x/y* direction was positioned to collect data from as many CA1 pyramidal cells as possible. Time-series images were collected through Scanbox (Neurolabware) and the PicoScope Oscilloscope was used to synchronize frame acquisition timing with behavior.

### Imaging sessions

The familiar environment was the same environment that the animals trained in. The experiment protocol for single day imaging sessions is shown in Fig. 1B. Each trial lasted ∼8-12 minutes and was always presented in the same order. Mice (n = 6) were first exposed to the familiar rewarded environment (Rewarded, R). Mice on average ran 34 ± 2 (mean ± standard error of the mean (s.e.m)) laps during this time, at which point, reward was turned off and imaging in the Unrewarded environment (UR) continued (30 ± 4 laps). In the Unrewarded condition, both reward and auditory cue associated with the reward (solenoid click) were disabled. Reward was then turned on again (Re-rewarded, RR) and mice ran 27 ± 3 laps. The mice were then introduced to a Novel-rewarded environment (NovR; 31 ± 5 laps). The Novel-rewarded environment (NovR) had distinct visual cues, colors and visual textures (Fig. 1B), but the same dimensions (2 m linear track) and reward location (end of the track) as the familiar environment. To rule out the possibility that observed changes in population activity were due to time, mice were exposed to only the familiar Rewarded environment for 20 minutes (control, n = 6).

The experiment timeline for multi day imaging sessions is shown in Fig. 3A. On Day 1, mice (n = 5) were exposed only to R. At the end of the imaging session, a 1 minute time-series movie was collected at a higher magnification and then averaged to aid as a reference frame in finding the same imaging plane on subsequent days (Fig. 3A). On Day 2, mice first experienced R followed by UR. On Day 3, UR was experienced first followed by RR. Where applicable, data from Day 2 was assessed together with data from the single day imaging sessions (combined n = 11 mice that were exposed to R followed by UR).

### Image Processing and ROI selection

Time-series images were preprocessed using Suite2p(Pachitariu et al., 2017). Movement artifacts were removed using rigid and non-rigid transformations and assessed to ensure absence of drifts in the *z*-direction. Datasets with visible *z*-drifts were discarded (n = 2). For multi day datasets, imaging planes acquired from each day were first motion corrected separately. ImageJ (NIH) was then used to align the motion corrected images relative to each other by correcting for any rotational displacements. The images across all days were then stitched together and motion corrected again as a single movie. Regions of interest (ROIs) were also defined using Suite2p (Fig. 1Aiii) and manually inspected for accuracy. Baseline corrected ΔF/F traces across time were then generated for each ROI and filtered for significant calcium transients, as previously described(Dombeck et al., 2010; Sheffield and Dombeck, 2015; Sheffield et al., 2017). Finally, we used raster plots(Stringer and Pachitariu, 2019) to visualize the ΔF/F population activity of neurons across time and across all conditions (Fig. 1C, Supplementary Fig. 3, Supplementary Fig. 4 and Fig. 3B). In these raster plots, neurons were clustered and sorted such that neurons with correlated activity were next to each other on the vertical axis (https://github.com/MouseLand/rastermap). For visual clarity, only neurons with at least 2 transients above 10% ΔF/F over the time of the experiment were included in the raster plot and the 2-D plots were interpolated using a hanning filter.

### Position Decoding

We trained a naive Bayes decoder (scikit-learn, Python(Pedregosa et al., 2012)) to predict the spatial location of the animal on the linear track from population activity within each mouse. Population activity consisted of ΔF/F traces from all identified cells organized as NxT, where N is number of cells and T is the total number of frames from an imaging session. Each lap traversal on the 2 m track was discretized into 40 spatial bins (each 5 cm wide). Time periods where the animal was stationary were filtered out (speed < 1 cm/s) and the decoder was only trained on frames belonging to running periods > 1 cm/s. Running behavior and population activity before and after filtering is shown in Supplementary Fig. 3 and Fig. 1C, respectively. To ensure decoder performance was not confounded by teleportation, we considered the end of the track as continuous with the beginning of the track so that the topology of the track was treated as a circle.

To assess how well a decoder trained in R was able to decode the animal’s spatial location in other conditions (Fig. 1 D-I), the decoder was trained on the first 60% of laps in R. The resulting model was evaluated on the remaining laps in R and on all laps in UR and RR (Fig. 1D). Decoder performance was assessed by calculating the coefficient of determination (R^2^) between the actual location of the animal and the location predicted by the decoder. Decoder error was quantified as the absolute difference in actual and decoded position in cm (Fig. 1G and H, 2C and D). We also trained and tested decoders within each condition in each mouse (Fig. 2A-E). Here, to assess decoder performance and to account for population activity changes across time, we employed a cross-validation approach by sliding the tested laps (20% of laps) by one each time and training on the remaining laps (80% of laps). To account for different numbers of laps across conditions, we down sampled each condition to match the condition with the least number of laps. To evaluate the influence of the number of cells on decoder performance, n% of cells from the total number of active cells detected in each mouse were randomly chosen and decoder performance was cross-validated. This was repeated 50 times and the mean R^2^ is reported in Fig. 2A.

### Decoder performance with different behavioral parameters

#### Licking Behavior

Licking data was collected using a capacitive sensor on the waterspout. Well trained mice showed a higher proportion of licks (pre-licking) in the region immediately preceding the reward in R (Supplementary Fig. 1). This anticipatory licking behavior continued for a few laps in UR (5 ± 1 lap) and decayed exponentially (Fig. 1E) with the exception of some animals (4/11) that randomly licked in later laps. To analyze the relationship between decoder error and licking, we identified the lap when licking had stopped in UR when 2 consecutive laps had no licks, and then divided the data into laps before licking stopped (UR, before licking stops) and laps after licking stopped (UR, after licking stops). We found that if we instead used a different criterion to identify when licking had stopped, i.e. the 1^st^ lap with no licks, or 4 consecutive laps with no licks, or 6 consecutive laps with no licks, our results were unaffected (Supplementary Fig. 5). This was also true if instead of defining when licking stopped in UR we simply grouped laps together based on the presence or absence of licks (Supplementary Fig. 6). From Fig. 1G onwards, UR refers to laps after licking stopped defined by our criteria of 2 consecutive laps of no licks. We obtained the lap wise decoder fit (R^2^) and lick frequency in UR in each animal and ran a rolling average with a sliding window of 3 laps (Fig. 1E). The average decoder fit across laps formed an S-shaped curve. We fit this mean R^2^ to a reverse Boltzmann Sigmoid curve (scipy.curve_fit, Python, Fig.1E, coefficient of determination of curve fit with mean decoder = 0.94). To calculate the inflection point at which the rate of decrease in R^2^ reaches the maximum, we calculated the first point where the second derivative of the fit reached 0 (lap 10, Fig 1E).

#### Time taken to complete a lap

This was calculated as the total time (in seconds) taken by the animal to run from 0 to 200 cm. We assessed if there was any correlation between the decoder fit and the time the animal took to complete a lap. To do so, we created a histogram of the distribution of time taken to complete a lap in R and UR (Supplementary Fig. 2). For each animal, we divided the laps in UR into those that overlapped with the histogram in R (Matched laps) and those that didn’t (Slower laps). The average time taken to complete a lap in the matched laps was 7.11 ± 0.25 s in R and 7.53 ± 0.20 s in UR. The slower speed laps took 21.67 ± 1.00 s. The majority of the laps belonged to the Matched speed laps and consisted of 71% of the total laps run by all animals in UR. Results are shown in Supplementary Fig. 2.

#### Attention

In R, mice slowed down as they approached the end of the track. We postulated that if mice were continuing to pay attention to where they were in VR when reward was removed, they would display a similar approach behavior. As a control, we first recorded running behavior of trained animals in the dark (n = 6), without any visual cues, to ensure that well trained mice were not displaying a stereotypical behavior independent from VR. To assess approach behavior, instantaneous velocity was calculated at each point along the 2 m track. This velocity trace was then smoothed by averaging it over 5 cm bins. In the dark, there were no signs of stereotyped behavior that looked like approach behavior (Supplementary Fig. 8). However, UR mice displayed the same approach behavior as in R on average (attended laps accounted for 88% of total laps in UR). We did find a larger variability in individual laps in UR, but when we removed these “unattended” laps from our analysis our decoder results remained unchanged.

### Defining Place Fields

Place fields were identified as described in previous studies(Dombeck et al., 2010; Sheffield and Dombeck, 2015; Sheffield et al., 2017) with a few key differences. The 2 m track was divided into 40 position bins (each 5 cm wide). The running behavior of the animal was filtered to exclude time periods where the animal was immobile (speed <1 cm/s). Filtering was done to ensure that place cells were defined only during active exploration. In UR, only frames after the licking stopped (see section on Licking Behavior) were included for place cell analysis. Place fields across the entire track were extracted as long as they began firing on the track. Cells that began firing at or after reward delivery and during teleportation were excluded from this analysis (although see Reward Cells below). Extracted place fields satisfied the following criteria and the same criteria was used for all conditions and all mice: 1. Their width was > 10 cm (with the exception of fields that are clipped at the end of the track). 2. The average ΔF/F was greater than 10% above the baseline. 3. The average ΔF/F within the field was > 4 times the mean ΔF/F outside the field. 4. The cell displayed calcium transients in the field on > 30% of laps. 5. The rising phase of the mean transient was located on the track. 6. Their p-value from bootstrapping was <0.05(Dombeck et al., 2010). Multiple place fields within the same cell were treated independently.

### Place Field Parameters

To calculate the various place field parameters, we binned the track into 40 bins (5 cm wide) and measured the mean ΔF/F of each bin. The data of each place field was a Lx40 matrix where L is the number of laps traversed by the animal. For all measures other than out-of-field firing and spatial correlation, transients outside the defined place field region were removed.

#### Center of mass

The COM from all traversals L was calculated as described in(Sheffield and Dombeck, 2015).

#### Reliability

Reliability of a place cell is the consistency with which it fires at the same location across multiple lap traversals. To calculate this, we computed the Pearson correlation between each lap traversal to obtain an LxL matrix. To obtain the reliability index, the average of this correlation matrix was multiplied by the ratio of number of laps with a significant calcium transient within the field and the total number of laps. The reliability index is 1.0 if the cell fires at the same location in each lap and 0.5 if it fires at the same position but only in half the laps, and so on.

#### Out/in place field firing ratio

This was computed as the ratio between the mean ΔF/F in bins outside the place field and the mean firing in bins within the place field.

#### Width

Width of the place field was computed as the distance between the spatial bin at which the mean place field rose above 0 and the spatial bin when it decayed back to 0. For place fields at the end of the track that were clipped the end of the place field was considered as the end of the track.

#### Firing Intensity

Firing intensity of the place field was calculated as the peak ΔF/F of the mean place field.

#### Spatial Correlation with Rewarded condition

To calculate the consistency of firing of the place cells defined in R across different conditions, we calculated the Pearson correlation coefficient between mean place cell activity defined in R and the mean of the Lx40 matrix of the same cells in other conditions. The within-session correlation was calculated from control animals (n = 6). The control rewarded condition (the duration control mice were in this condition matched experimental mice that experienced R-UR) was divided into two halves and the correlation coefficient was calculated between the mean place cell firing in the two halves.

#### Spatial correlation across days

Place cells were defined on Day 1 in R or UR conditions. Mean place field of each identified cell was then correlated with the mean of the Lx40 matrix on Day 2. Stable cells were defined as those cells with a correlation coefficient > 0.5. This was chosen as a threshold based on correlations observed within the same day: 98% of cell correlations observed within the same day were above 0.5.

### Network Analysis

Network analysis was performed on place cells. Place cells (N) were defined on Day 1 in R or UR. Pairwise Pearson’s correlation coefficient was calculated between mean place field activity on Day 1 and mean place field activity on Day 2 to form a NxN adjacency matrix. We also calculated an adjacency matrix from correlation coefficients between two halves of the Day 1 session in R (Control), and between Day 1 in R and shuffled place cell activity on Day 2 (shuffle). Correlation coefficients that were negative and not statistically significant were removed. Networks were plotted using the open-source tool Gephi (https://gephi.org/). Place cells were nodes and weights of the edges between two nodes were the correlation coefficients. We used the Fructherman Reingold layout as it is a force directed layout algorithm that prioritizes placing nodes connected by edges with high weights close together. In a highly stable network, where place cells are only correlated to themselves and other place cells with similar fields, nodes are tightly clustered by place field location, resulting in a spring-like topology (Fig. 3I; Left: within session Control). A similar topology was evident across days in R, but not in UR or in the shuffled dataset. We calculated the weighted degree and cluster coefficient for each node using built-in functions in Gephi. On the layout in Fig. 3I, nodes were sized by their weighted degree (higher the weighted degree, larger the circle) and colored by their place field position on Day 1. To display weighted degree and cluster coefficients across track position, we binned nodes by their center of mass on the track, each bin being 25 cm in length.

### Reward Overrepresentation

To compute the density of place cells along the track, the COM of all place fields in all animals were fitted to a gaussian distribution (mean ± standard deviation of the gaussian distribution in cm in different conditions, R: 114 ± 55, UR: 102 ± 54, RR: 112 ± 58, NovR: 108 ± 53) and a uniform distribution to extract regions of place cell overrepresentation (Supplementary Fig. 9). The excess field density in R above the uniform distribution was at 175-200 cm (25 cm before the reward zone). We defined this 25 cm region as the end of the track. To compare changes in place field density across conditions between the middle of the track (25-175 cm) and end of the track (175-200 cm), we divided the middle of the track into 25 cm bins and averaged place cell density across the bins (Supplementary Fig. 9).

### Reward cells

Reward cells were defined as described in(Gauthier and Tank, 2018). A cell was defined as a reward cell if it fired at the reward zone on the track (40 cm before reward) and around reward delivery (2 seconds before and after reward delivery) in both R and the NR. The reward zone on the track was chosen based on the area of high place field density before the reward in R and N (Supplementary Fig. 11). In total, we found 43 such cells from 6 animals, both on track and around reward delivery (Supplementary Fig. 11A). These cells constituted 0.9 % of all active cells recorded. To compare reward cell firing across all conditions, we computed the lap wise firing of these cells in time around reward delivery. Their COM in time around reward delivery, reliability and correlation with R was then calculated similar to place cells.

### Statistics

For data distributions, a Shapiro-Wilk test was performed to verify if the data was normally distributed. If normality were true, where applicable, a paired or unpaired Student t-test was used. For non-normal distributions, a paired Wilcoxon signed rank test or an unpaired Mann-Whitney U test was used. For samples with five data points or less, only a non-parametric test was used. Multiple comparisons were corrected using one-way ANOVA with Bonferroni post-hoc. Box and whisker plots were used to display data distributions where applicable. The box in the box and whisker plots represent the first quartile (25^th^ percentile) to the third quartile (75^th^ percentile) of the distribution, showing the interquartile range (IQR) of the distribution. The black line across the box is the median (50^th^ percentile) of the data distribution. The whiskers extend to 1.5*IQR on either side of the box. A data point was considered an outlier if it was outside the whiskers or 1.5*IQR. Significance tests were performed with and without outliers. Data distributions were considered statistically significant only if they passed significance (p < 0.05) with and without outliers. To model the probability distribution in the datasets and get an accurate idea of the data shape, a kernel density estimate was fitted to the data distribution and is shown alongside histograms. Cumulative probability distribution functions were compared using a Kolmogrov-Smirnov test. We employed estimation statistics to ascertain the level of differences between distributions by using the DABEST (Data Analysis with Bootstrap-coupled Estimation) package(Ho et al., 2019). Estimation plots display the median difference between two conditions against zero difference, with error bars displaying 95% confidence intervals of a bootstrap generated difference (5000 resamples). A kernel density fit (shaded curve) on the resampled difference is also displayed alongside. This difference was compared against zero. Correlations were performed using Pearson’s correlation coefficient. p < 0.05 was chosen to indicate statistical significance and p-values presented in figures are as follows: *, p < 0.05, **, p < 0.01, ***, p < 0.001, N.S. not significant. Data summary in the text and error bar and shading in figures are presented as mean ± s.e.m, unless stated otherwise. Data preprocessing and place field extraction was done on MATLAB (Mathworks, Version R2018a). All other data and statistical analyses were conducted, and figures were made, in Python 3.7.4 (https://www.python.org/).

## Acknowledgments

We thank M. Howe, J. Heys, and N. Spruston for comments on the manuscript, D. Freedman for comments on the Cover Letter, M. Rosen for assistance on data analysis, C. Heer for discussions on dopaminergic inputs, H. Macomber for behavior data of mice running in the dark and members of Sheffield lab for manuscript comments and useful discussions.

## Author contributions

C.C. performed surgeries. C.C and S.K. collected the data. S.K. wrote the analysis code and analyzed data. S.K. and M.S. conceived and designed the experiments, interpreted the data and wrote the manuscript.

---

**Supplementary Fig. 1.**
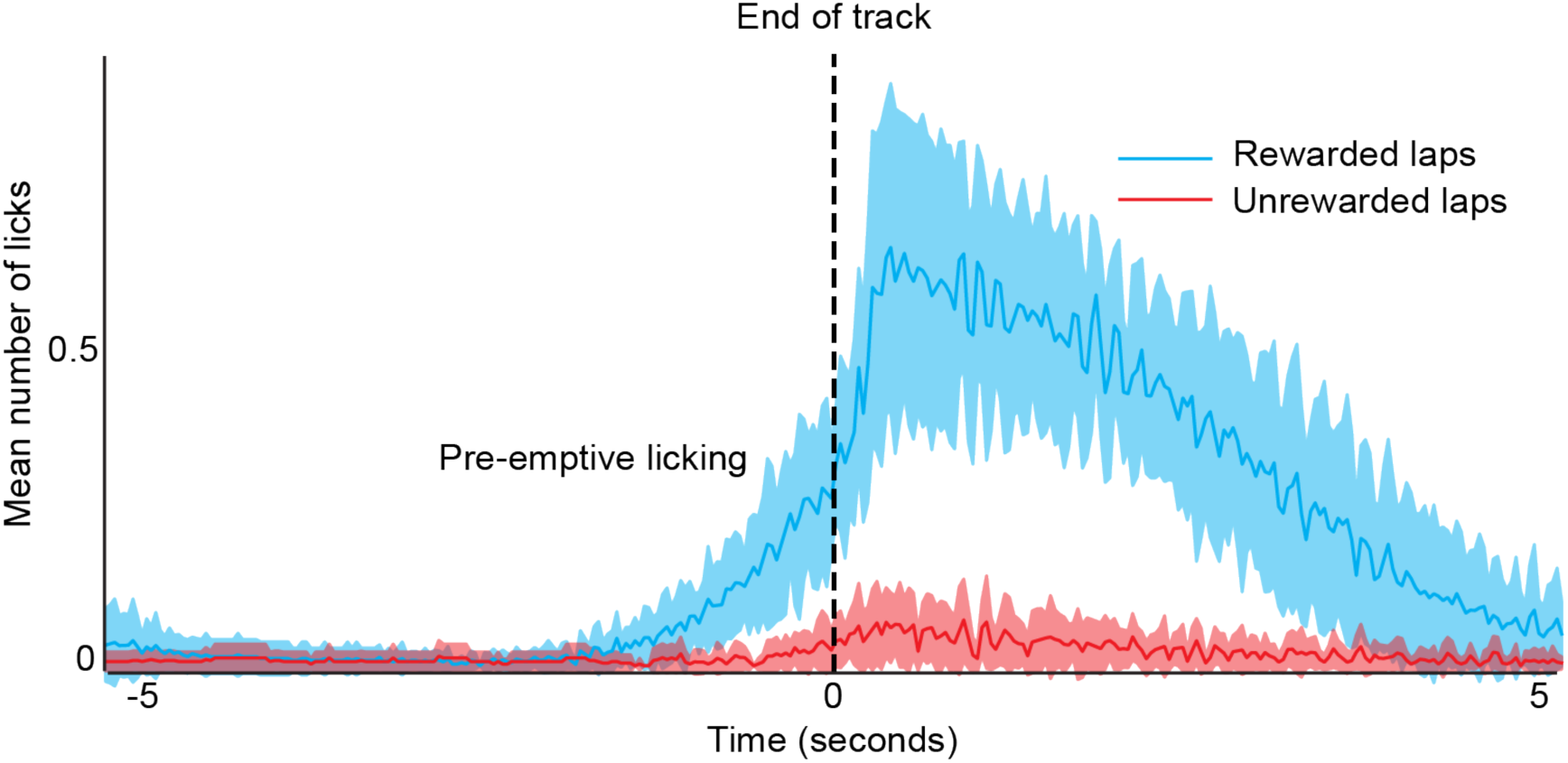
Mice pre-emptively lick before a learned reward location in VR environment. Mean number of licks around reward delivery (time = 0) in Rewarded (blue, R) and Unrewarded condition (red, UR). Number of licks were calculated for each mouse on each lap and were binarized as 1 or 0 depending on if the animal licked or not in the given time bin. In R, animals display preemptive licking in anticipation of the reward, which was absent in UR after a few laps. Shading represents standard deviation.

**Supplementary Fig. 2.**
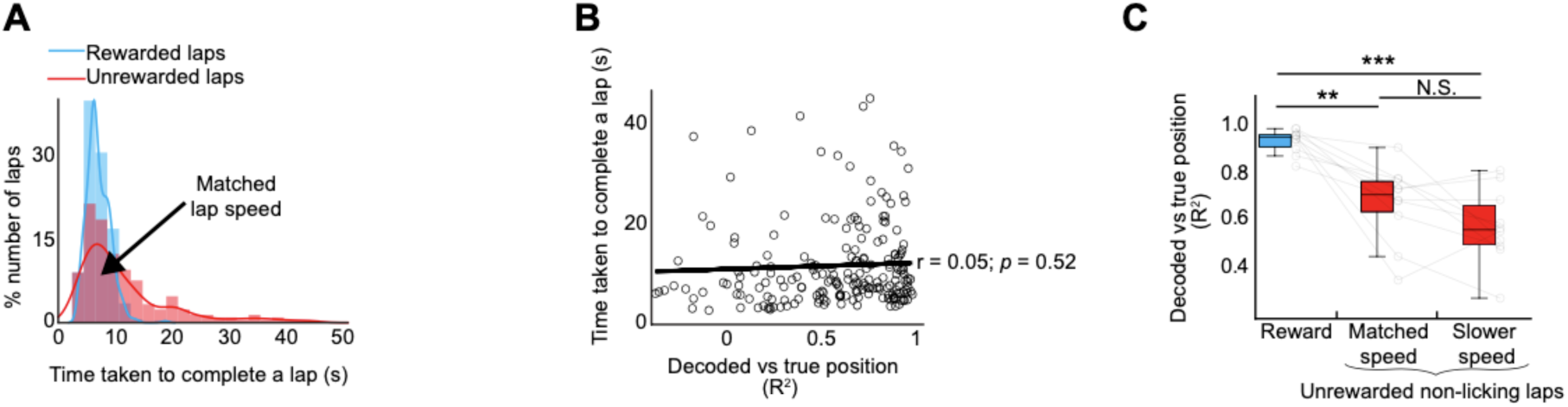
Diminished decoder performance in the unrewarded condition is not due to changes in running behavior. (A) Histogram of lap speed defined as the time taken to complete a lap (in seconds) on the 200 cm track in R and UR (n = 11 mice, total laps in R = 409, UR = 324). Curves are kernel density estimates on the distribution. 71% of the laps in UR had lap speeds that matched lap speeds in R (matched speed laps = 229, slower speed laps = 95). (B) Scatter plot between lap speed and decoder performance (coefficient of determination between true position and decoded position, R^2^). Each circle is a lap (data pooled from all mice). Linear regression (y = 11 + 0.9 * x) is shown in black, r and p-value were derived from Pearson’s correlation coefficient. (C) Mean R^2^ for each mouse (circles). Laps in UR were separated into laps with speeds that matched laps in R and laps that were slower. The box and whisker plots show the distribution of the mean R^2^ for each condition. P-values were obtained using a Paired t-test, ANOVA with Bonferroni post hoc was done to correct for multiple comparisons. P-values were obtained with and without outliers. If P-values were significant in both situations, then the P-value with outliers is displayed here. **P < 0.01, ***P < 0.001.

**Supplementary Fig. 3.**
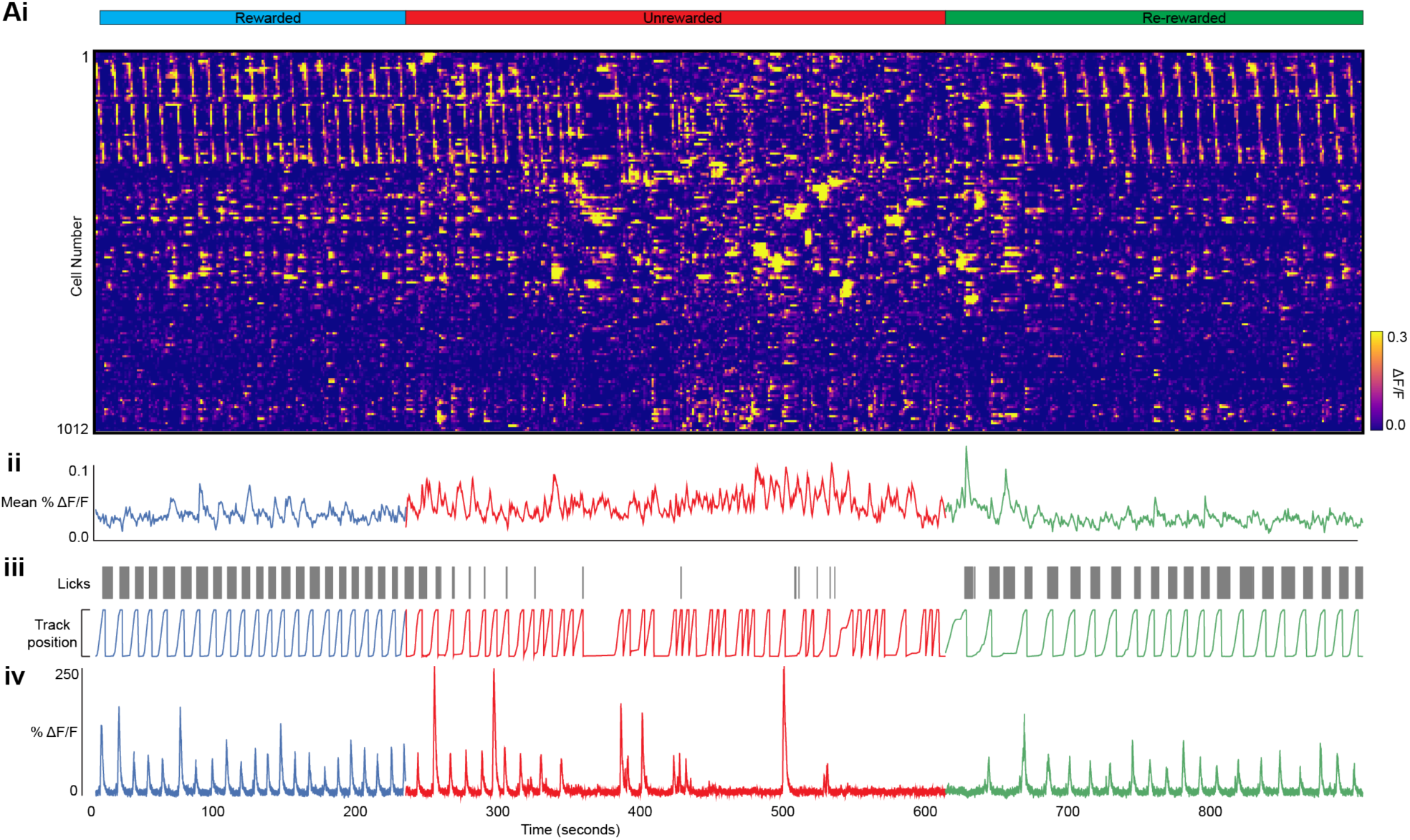
Population activity without filtering laps for good running behavior. This is the same dataset as Fig. 1C but without removing periods where the mouse was stationary. Most laps look similar to the filtered version shown in Fig. 1C because mice run in VR consistently even when reward is removed. Most stationary periods are found at the start of the track before the mouse begins its traversal of the environment. Unfiltered laps are shown in (iii).

**Supplementary Fig. 4.**
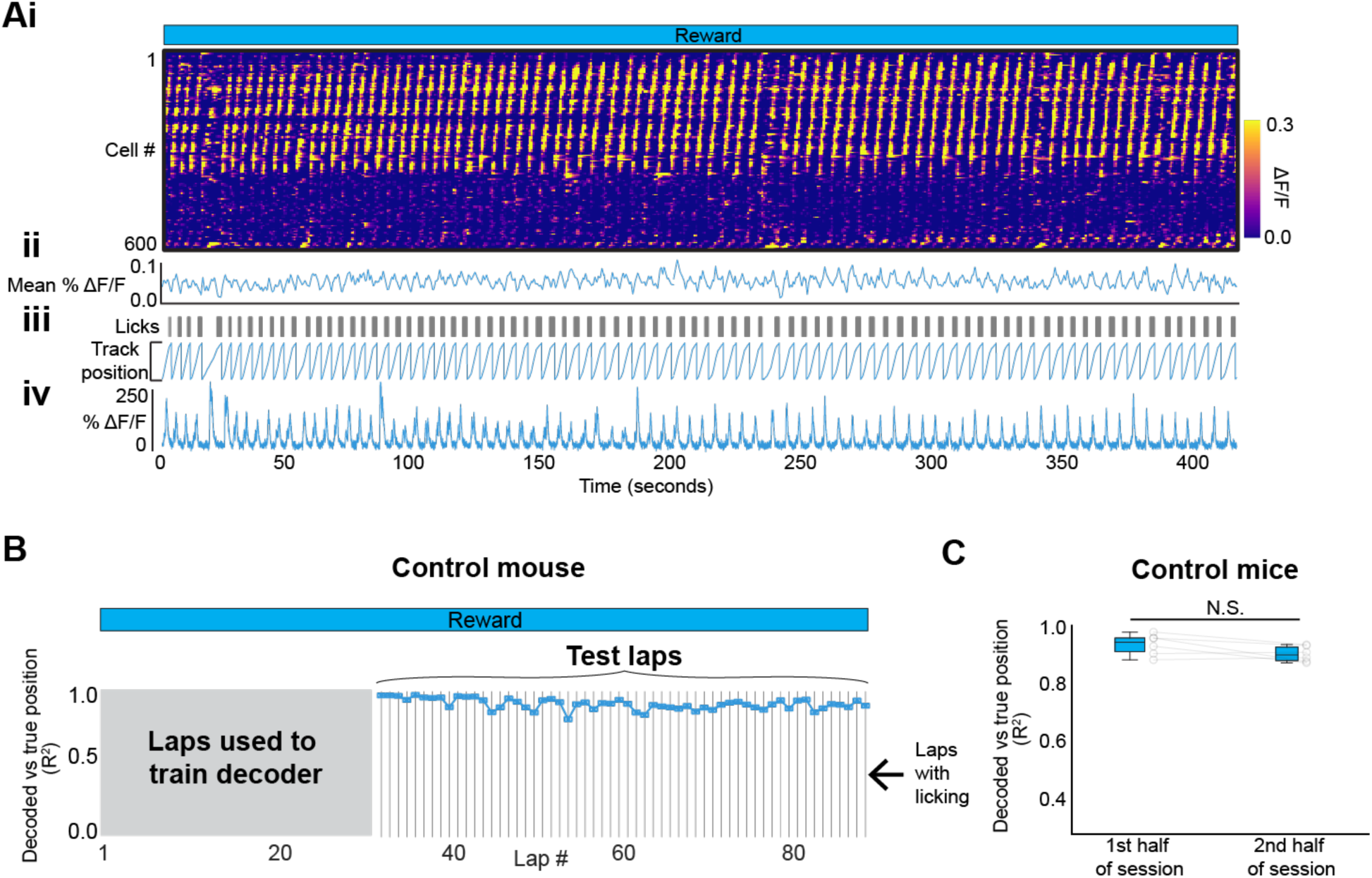
Population activity remains stable with time. (A) (i) Raster representation of population activity in 1 mouse traversing the rewarded environment for 15 minutes (a similar amount of time and number of laps over which experimental mice were switched between R and UR conditions). (ii) Mean activity of cells above (iii) mouse behavior (iv) example cell. Note the relative stability of the population activity compared to Fig. 1C. (B) Bayesian decoder trained on activity on initial 30 laps to predict animal’s position on the other laps. 30 laps were chosen as they matched the average number of laps used to train the decoder in the experimental rewarded condition (see Fig. 1D). Decoder performance (coefficient of determination, R^2^) between decoded and true position in the tested laps are shown. (C) Tested laps were divided into two halves and mean R^2^ of true vs predicted position for each mouse in each half of the session is displayed.

**Supplementary Fig. 5.**
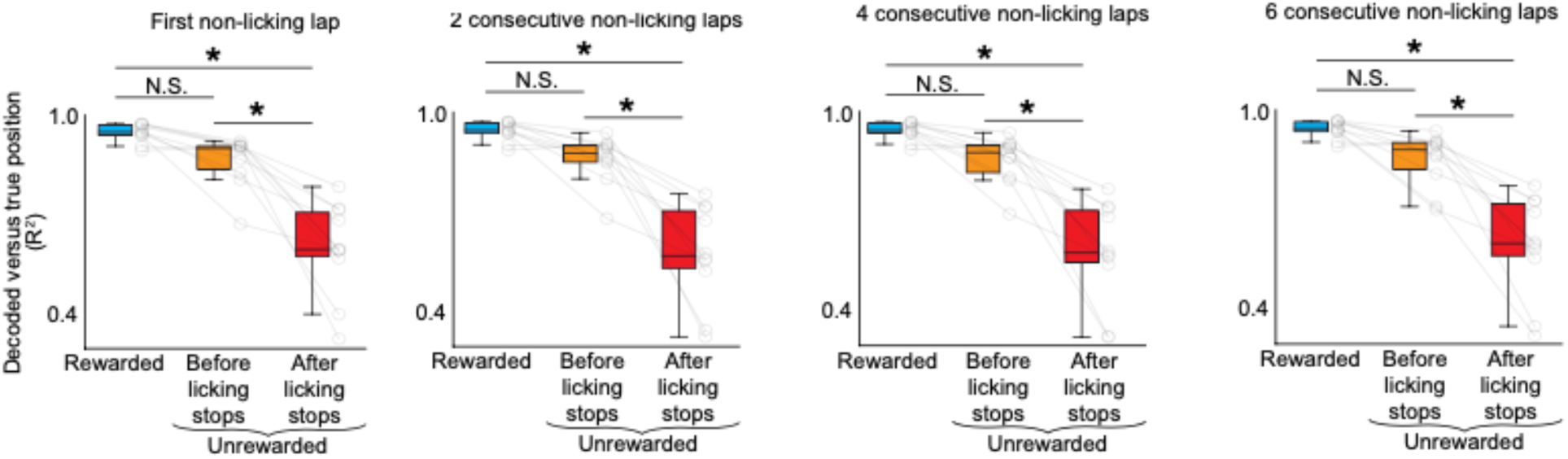
The change in decoder performance in the unrewarded condition after licking stops does not depend on our definition of when licking stops. Mean lap decoder performance (coefficient of determination, R^2^) between decoded and true position for each mouse (circles) in the tested laps in Rewarded and Unrewarded condition. Unrewarded laps are separated by before and after licking stops. The definition of the lap at which licking stops varies in each panel as indicated in the title. P-values were obtained using a Paired t-test. ANOVA with Bonferroni post-hoc was used for multiple comparison correction. *P < 0.05.

**Supplementary Fig. 6.**
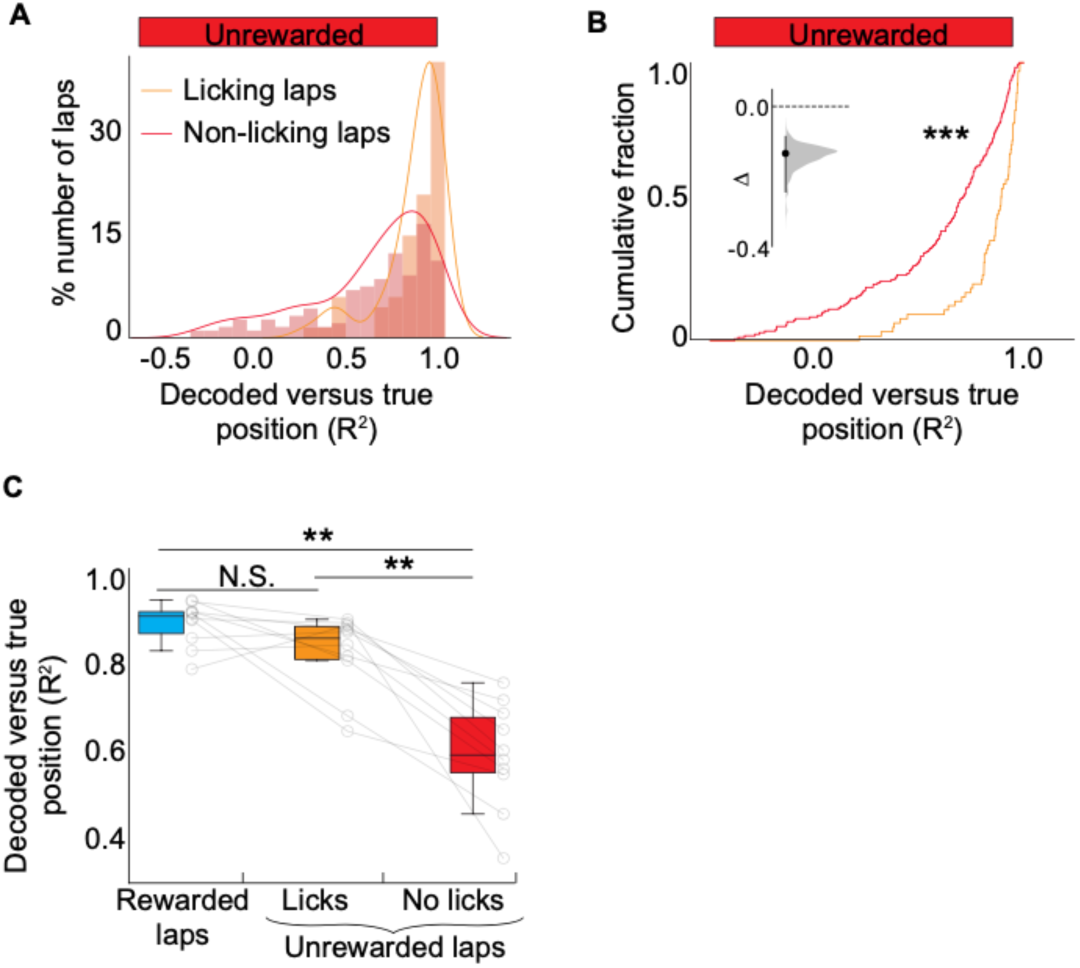
Decoder performance between laps with licks and laps without licks. (A) Histogram of decoder performance (R^2^) in all laps with licks and all laps without licks in the Unrewarded condition (instead of defining the lap in which licks stopped). Curves are a kernel density estimate on the distribution. (B) Cumulative distribution. P-values were obtained using KS-Test. Inset shows an estimation plot of median difference between the two distributions, compared against zero difference. Error bar is bootstrap (5000 resamples) generated 95% confidence intervals. Grey shaded area is a kernel density estimate on the resampled distribution. Plots were made using Data Analysis with Bootstrap-coupled Estimation (DABEST, (Ho et al. 2019)). (C) Mean decoder performance (R^2^) for each mouse (circles), laps in UR are separated into laps with licks and laps without licks. P-values were obtained using a Paired t-test, ANOVA with Bonferroni post hoc was done to correct for multiple comparisons. **P < 0.01.

**Supplementary Fig. 7.**
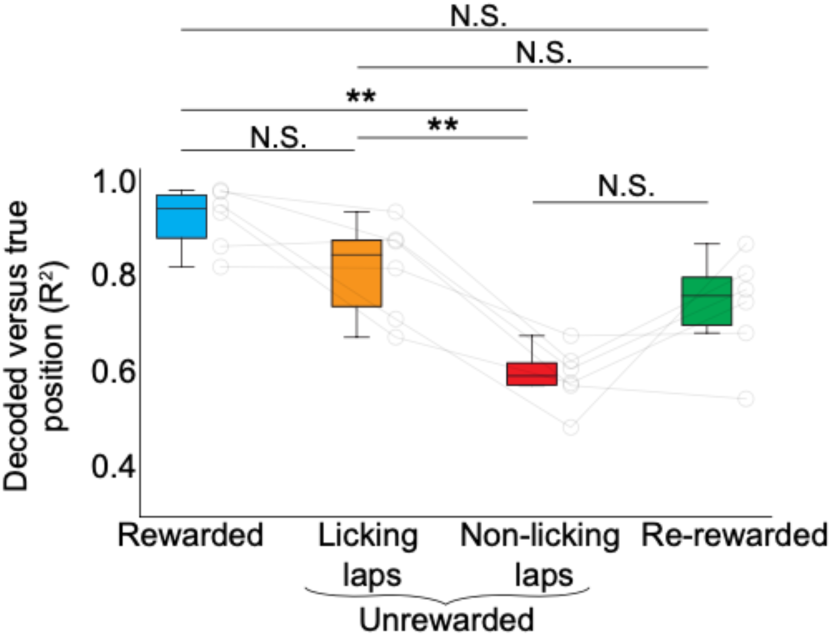
Decoder performance increases in the Re-rewarded condition but does not return to Rewarded condition levels. Mean decoder performance (R^2^) on tested laps per animal (circles, n = 6) in Rewarded, Unrewarded and Re-rewarded conditions. Unrewarded laps were divided into laps with licks and laps without licks. P-values were obtained using a Paired t-test, ANOVA with Bonferroni post hoc was done to correct for multiple comparisons. **P < 0.01.

**Supplementary Fig. 8.**
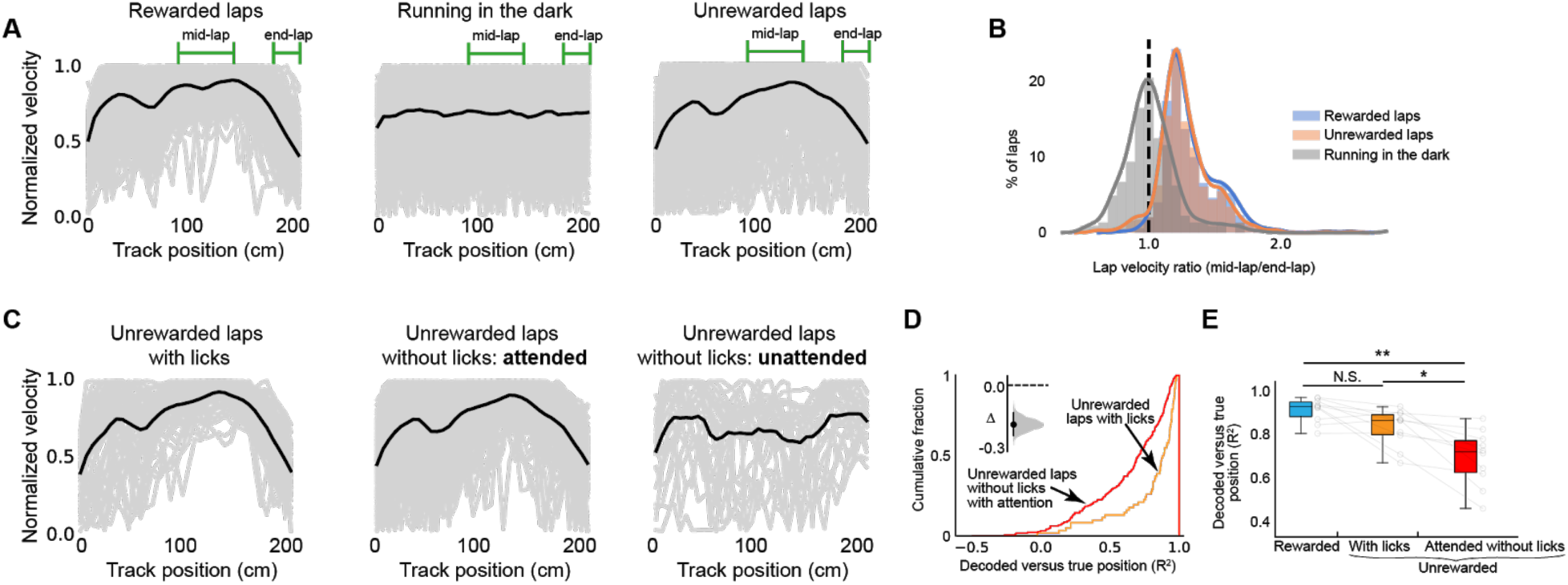
Diminished decoder performance in the Unrewarded condition is not due to inattentiveness to the VR environment. (A) Instantaneous velocity for each lap (gray traces) from 11 mice in Rewarded (n = 409 laps) and Unrewarded conditions (n = 324) and 6 mice in Dark condition (n = 226). The velocity on each lap was normalized to its peak. Mean velocity from all laps is shown in black for each condition. (B) Distribution of ratio between lap velocity in the middle (100-150 cm) and end (175-200 cm) of the track as indicated above the traces in each condition in A. Curves are kernel density estimates on the distributions. Laps were considered attended if the ratio was greater than 1 (dashed line). (C) Same as A but with laps in Unrewarded condition divided into laps with licks (n = 74; left), and laps without licks that were attended (n = 210; middle) and that were not attended (n = 40; right). (D) Cumulative distribution of decoder performance (R^2^) on unrewarded laps with licks (orange) and attended unrewarded laps without licks (red). Inset shows an estimation plot of bootstrapped median difference between the two distributions. P-values were obtained using KS-Test. Note the difference in decoder performance without licks was still lower than with licks even though mice were paying attention to their position on the VR track on these laps, indicating a drop in reward expectation causes this change and not a loss of attention. (E) Mean decoder performance (R^2^) in each mouse (circles). P-values were obtained using a Paired t-test, ANOVA with Bonferroni post hoc was done to correct for multiple comparisons. *P < 0.05, **P < 0.01, ***P < 0.001.

**Supplementary Fig. 9.**
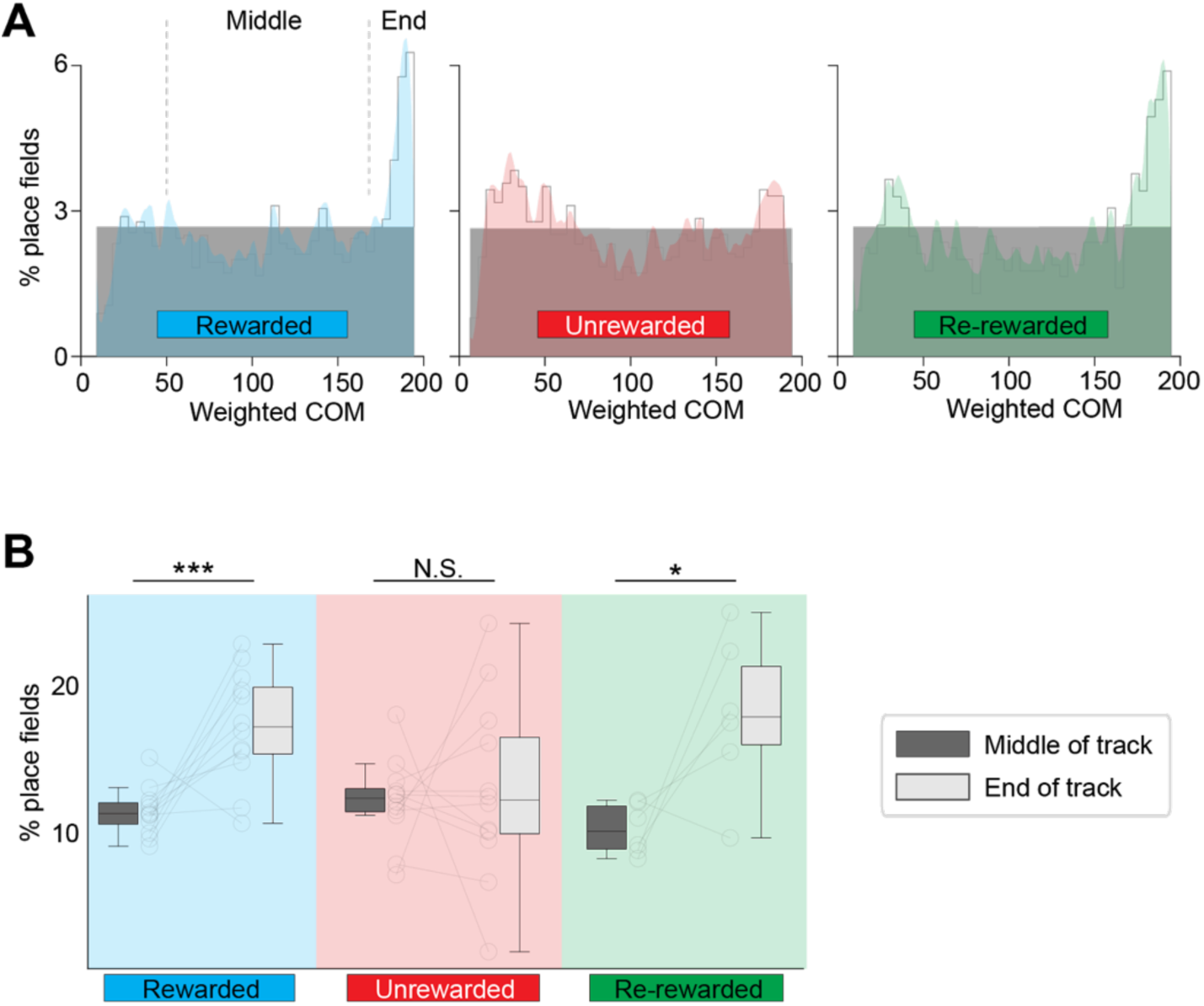
Over-representation of place fields near the reward site disappears in the unrewarded condition and reappears in re-rewarded condition. (A) Distribution of place field center of mass (COM) locations in each condition pooled from all mice. Plots show observed density (gray line), uniform distribution (gray shade) and Gaussian distribution (color). (B) Percentage of place fields in the middle of the track versus end of the track. *P < 0.05. ***P < 0.001. n = 11 mice in Rewarded and Unrewarded conditions, and n = 6 in Re-rewarded condition.

**Supplementary Fig. 10.**
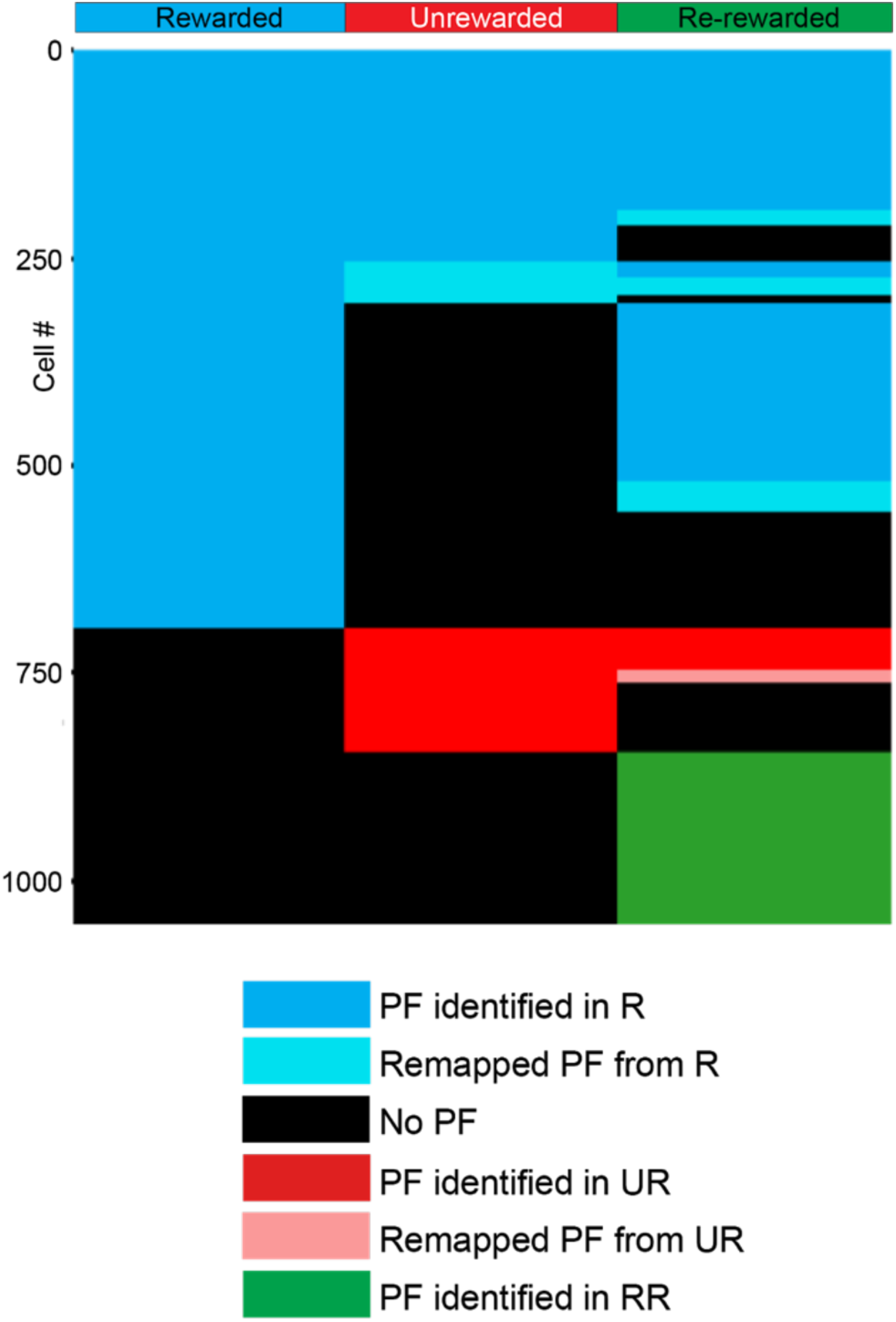
The formation and fate of place fields across conditions. Place fields identified in the rewarded condition (R, blue) can be stable throughout the Unrewarded (UR, blue) and Re-rewarded (RR, blue) conditions. They can also remap in UR or RR (cyan) or lose their place field completely in UR or RR (black). New place fields can form in UR (red) and be stable (red), remap (pink), or disappear in RR (black). New place fields can also form in RR (green).

**Supplementary Fig. 11.**
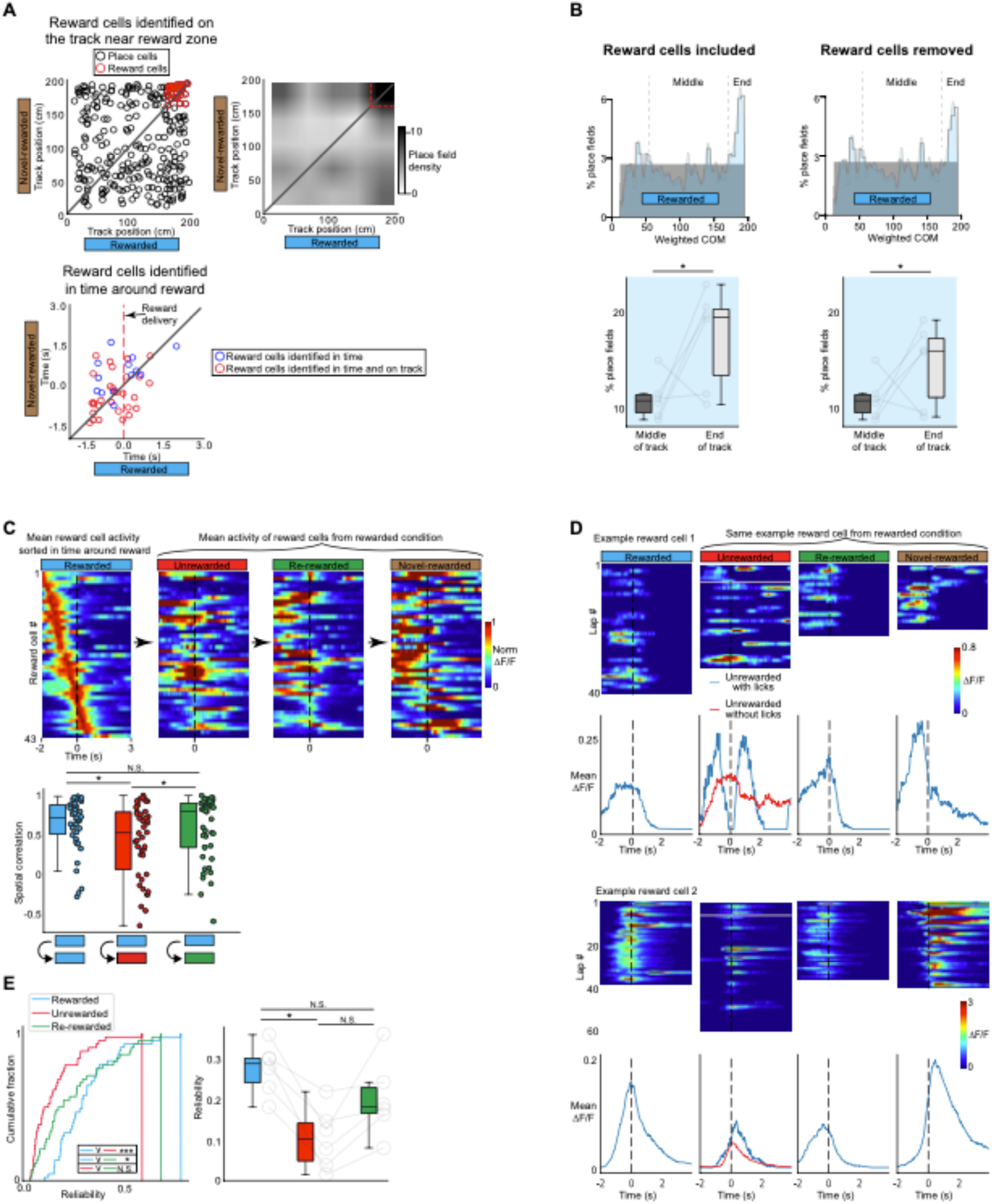
Reward cell encoding is disrupted in the unrewarded condition. (A) (Top left), center of mass (COM) of all cells with place fields (circles) in both the Rewarded (R) and Novel-rewarded (NovR) conditions from 6 mice (n = 273 cells). In both environments, reward was given at the end of the track and cells that fired around the reward were considered to be reward cells (30 cells, red circles). (Top right), density of COMs between the two conditions, spatially binned and smoothed (width 25 cm). Red lines are drawn around the defined reward zone based on increased place cell density (160 - 200 cm). (Bottom left), time of peak firing of cells from reward delivery. Cells that fired 1.5 seconds before and after reward delivery in both R and NovR were also identified as reward cells (13 cells). Peak firing time of the cells that were identified on the track (top) are colored in red. (B) (Top), distribution of place field center of mass (COM) locations in R with and without reward cells. Plots show observed density (gray line), uniform distribution (gray shade) and Gaussian distribution (color). (Bottom), percentage of place fields in the middle of the track versus end of the track in the two halves. P-values were obtained using a Paired t-test. (C) (Top), mean activity of identified reward cells from time of reward delivery (dashed line) in all conditions. Cells were sorted by their time of peak firing in R, and plotted in the same order in the other conditions. Cells were normalized to their peak firing in R. (Bottom), correlation coefficient of the same cells (dots) within R and between R and other conditions. P-values were obtained using a Paired t-test. (D) Example of two reward cells. Their lap by lap activity is shown on top and average activity at the bottom. Dashed line indicates the time of reward delivery. Pink line in the Unrewarded condition indicates the lap when the animal stops licking. (E) Trial-by-trial reliability of reward cells across conditions (See methods) as a cumulative distribution function (left, P-values: KS-Test) and per animal (dots, right). P-values for boxplot distributions in C and E were obtained using a Paired t-test, ANOVA with Bonferroni post hoc was done to correct for multiple comparisons. *P<0.05, **P<0.01 ***P<0.001.

**Supplementary Fig. 12.**
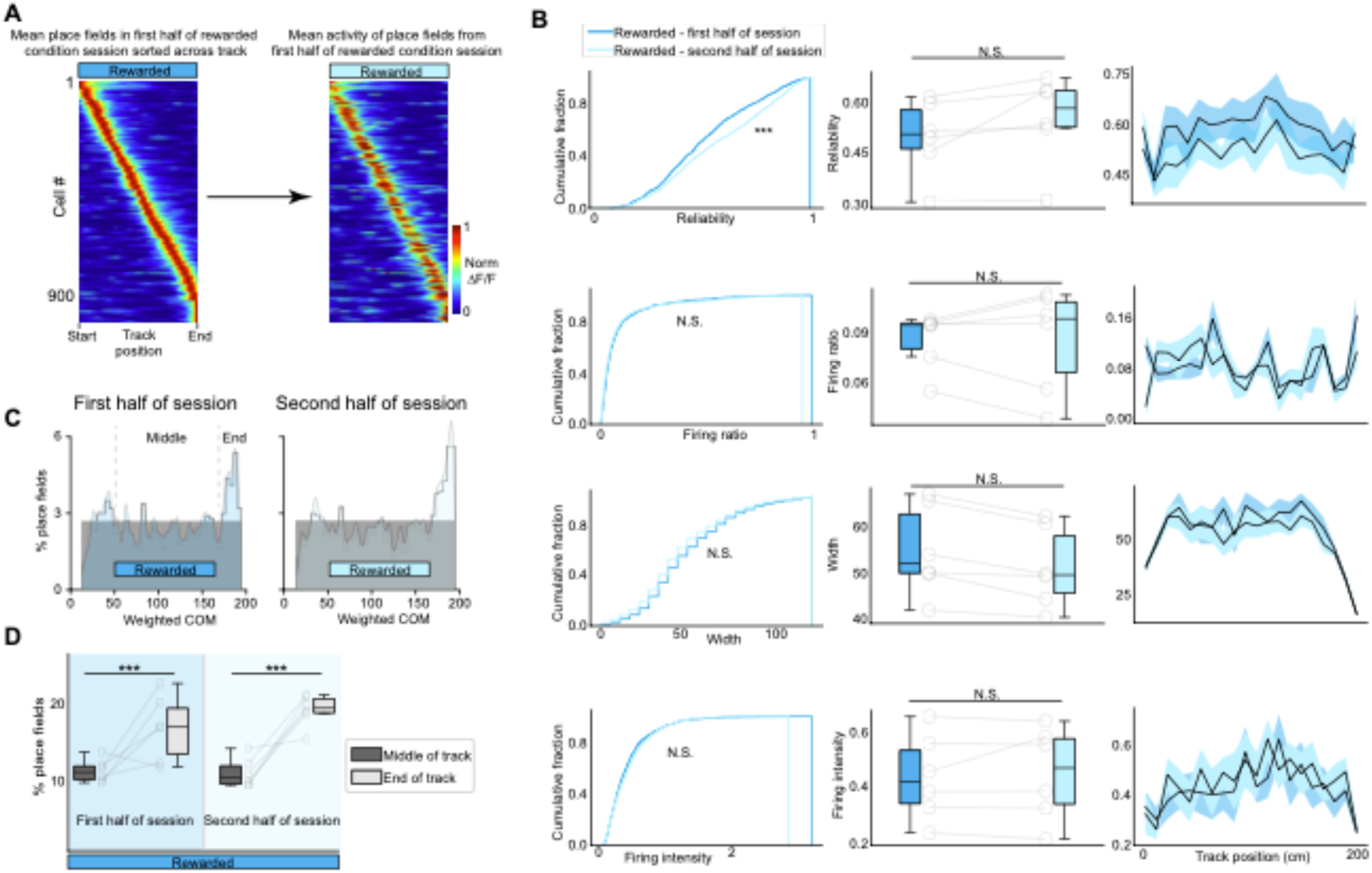
Place fields in control mice. Control mice (n = 5) were exposed only to the familiar rewarded condition for 15 minutes. The session was divided into two and place fields from the 2 halves were analyzed for changes in place field parameters. (A) Mean place fields of pooled place cells from all mice defined in the first half (left) and the second half (right). Place cells were sorted by their center of mass and normalized to their peak in the first half. (B) Place cell parameters pooled from all mice as a cumulative distribution function (CDF, left), means from each mouse (middle), and pooled from all mice by track position (right). See methods section for what these parameters are and how they were calculated. (C) Distribution of place field center of mass (COM) locations. Plots show observed density (gray line), uniform distribution (gray shade) and Gaussian distribution (color). Note the over-representation of place fields towards the end of the track in both cases. (D) Percentage of place fields in the middle of the track versus end of the track in the two halves. ***P < 0.001.

**Supplementary Fig. 13.**
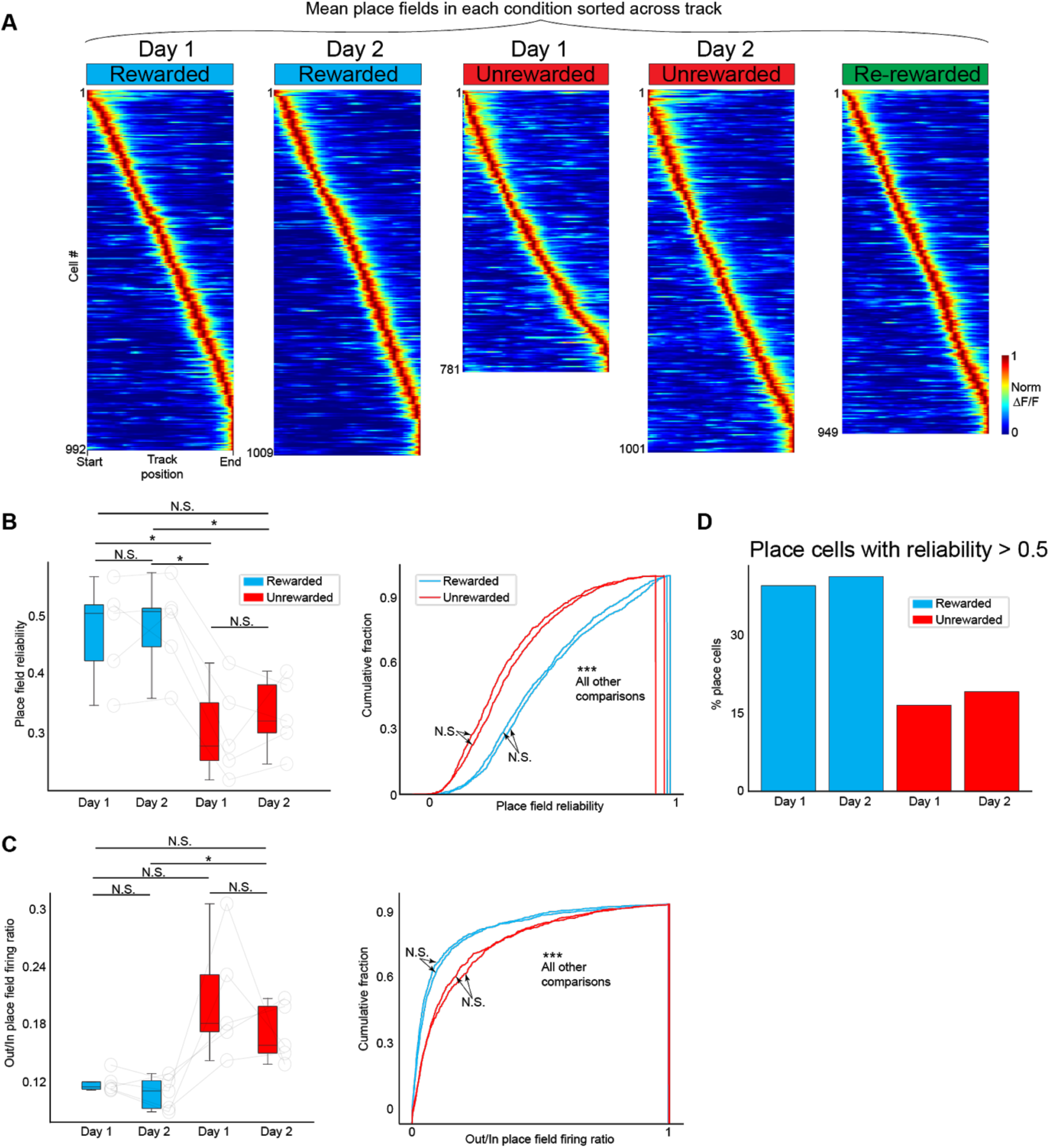
Place fields are more reliable across days in the rewarded versus unrewarded conditions. (A) Mean place fields determined in each condition, on each day, sorted by track position and pooled from all mice (n = 5). Activity of each cell is normalized to its peak. (B and C) Place field trial-by-trial reliability (B) and Out/In field firing ratio (C) in the rewarded and unrewarded conditions. (left) mean per animal (right) cumulative distribution function. (left), P-values were obtained using a paired Wilcoxon signed rank test and Kruskal Wallis with Bonferroni post hoc to correct for multiple comparisons. (right), P-values were obtained using KS-Test. (C) Percentage of place fields with reliability > 0.5 in each condition.

**Supplementary Fig. 14.**
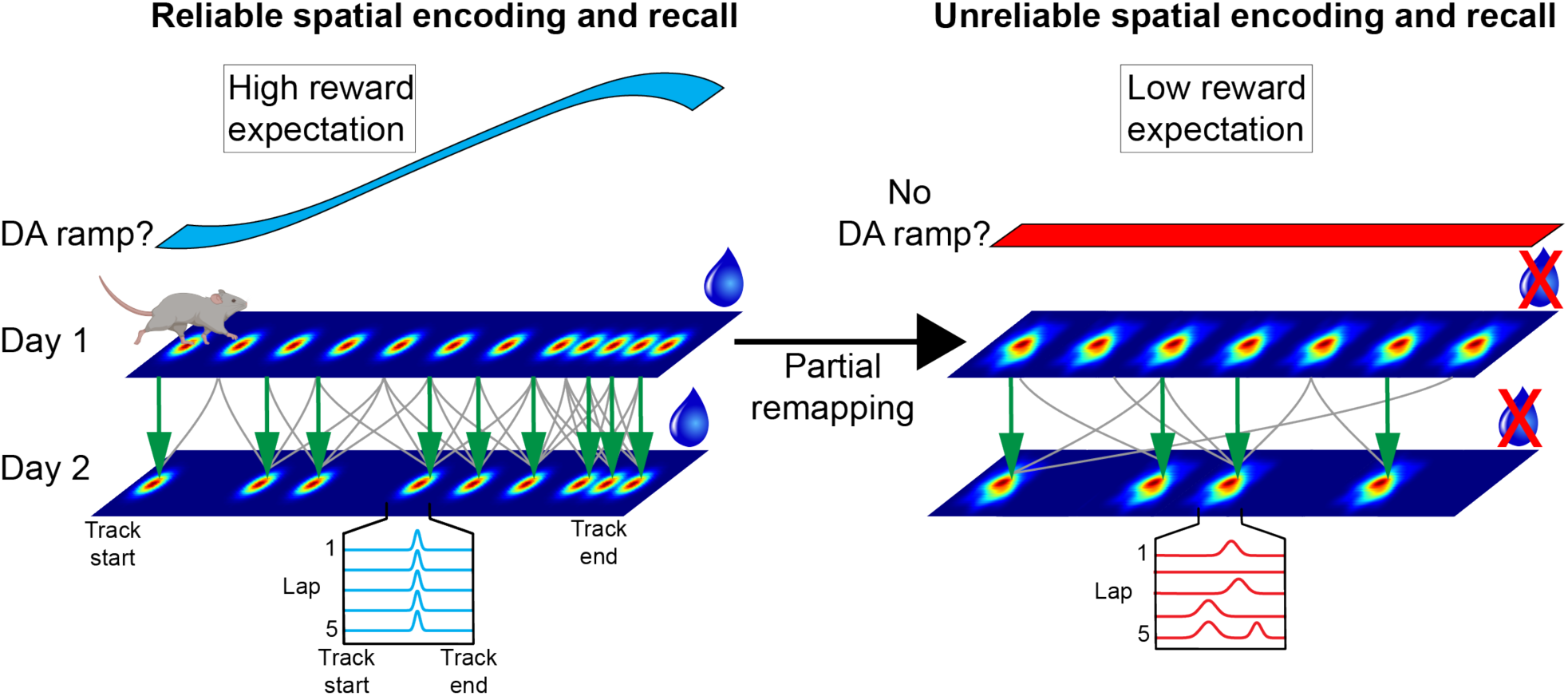
Conceptual model for how reward expectation influences spatial encoding and recall. In a familiar environment that has been consistently rewarded, mice develop a high reward expectation whenever they are in that environment (left). Under these conditions, place cells encode the environment reliably (example of lap-by-lap place field transients from a single place cell shown in blue at the bottom) with a precise map that over-represents locations near the reward site. Relative place field stability exists at the cellular (green arrows) and ensemble level (gray lines) over days.

In this scenario, we posit a possible mechanism where dopamine release from the ventral tegmental area (VTA) into the hippocampus ramps up as the animal approaches the rewarded location with high reward expectation (Mohebi et al. 2019; Howe et al. 2013; Hamid et al. 2016). Supporting this idea, we observed a ramping increase in the correlation of ensembles of place fields across days with proximity to reward that looks remarkably similar to ramping dopamine signals seen in other brain regions (Fig. 3M). Specifically, this dopamine ramp could promote glutamatergic synaptic transmission of specific sets of inputs that drive place cell firing in real-time, thus promoting lap-by-lap place field reliability (Tritsch and Sabatini 2012; Martig and Mizumori 2011). Dopamine could also facilitate long-term strengthening of the same synapses (Rosen et al. 2015; Ghanbarian and Motamedi 2013; Huang and Kandel 1995) to support the stability of place fields across days (Kentros et al. 2004). Although a dopamine ramp means the level of dopamine is not equal across the track, attractor-like dynamics could ensure dopamine influences place cells at all locations. On the basis of the known connectivity of CA1 neurons, this may arise from local inhibition within CA1 or driven by input from CA3, which does have recurrent connectivity to support attractor dynamics and also receives VTA input (Gonzalez et al. 2019; Lisman and Grace 2005; Colgin et al. 2010; Knierim and Zhang 2012; Rolls 2007; Gasbarri et al. 1994).

When reward expectation diminishes (right), the dopamine ramp disappears (Mohebi et al. 2019; Howe et al. 2013). This likely leads to rapid disruption of glutamatergic synaptic transmission (Tritsch and Sabatini 2012) triggering partial remapping of the place cells and causing the new map to become unreliable at all locations in the environment (Mamad et al. 2017; Gill and Mizumori 2006; Martig and Mizumori 2011). The loss of dopamine would also result in a loss of facilitation of synaptic potentiation, leading to reduced place field stability across days (Rosen et al. 2015; McNamara et al. 2014; Retailleau and Morris 2018; Martig and Mizumori 2011).

